# A midbrain - thalamus - cortex circuit reorganizes cortical dynamics to initiate planned movement

**DOI:** 10.1101/2020.12.16.423127

**Authors:** Hidehiko K. Inagaki, Susu Chen, Margreet C. Ridder, Pankaj Sah, Nuo Li, Zidan Yang, Hana Hasanbegovic, Zhenyu Gao, Charles R. Gerfen, Karel Svoboda

## Abstract

Motor behaviors are often planned long before execution, but only released after specific sensory events. Planning and execution are each associated with distinct patterns of motor cortex activity. Key questions are how these dynamic activity patterns are generated and how they relate to behavior. Here we investigate the multi-regional neural circuits that link an auditory ‘go cue’ and the transition from planning to execution of directional licking. Ascending glutamatergic neurons in the midbrain reticular and pedunculopontine nuclei show short-latency and phasic changes in spike rate that are selective for the go cue. This signal is transmitted via the thalamus to the motor cortex, where it triggers a rapid reorganization of motor cortex state from planning-related activity to a motor command, which in turn drives appropriate movement. Our studies show how brainstem structures can control cortical dynamics via the thalamus for rapid and precise motor behavior.

## Introduction

Many behaviors, including purposeful movements, are composed of distinct phases that require different computations. For example, while waiting at a red light to make a turn, we plan to rotate the steering wheel while pressing the gas pedal, and execute this motor program when the signal turns green. The planning and execution phases are produced by distinct patterns of neuronal activity (Svoboda and Li, 2018; Vyas et al., 2020). To support adaptive behavior, behavior-related neural activity rapidly switches from one pattern to another at the appropriate time, often guided by contextual cues, as in many laboratory decision-making tasks (Funahashi et al., 1989; Sommer and Wurtz, 2002; Kaufman et al., 2016, 2014; Vyas et al., 2020). The multi-regional pathways and mechanisms underlying these dynamic cortical activity patterns are not understood.

Here we track the multi-regional neural signals that link contextual cues and the switch from movement planning to movement initiation. Planned movements that are released by a contextual “Go cue” are faster and more precise than unplanned movements (Hanes and Schall, 1996; Rosenbaum, 1980; Shenoy et al., 2013). Planned movements are anticipated by slowly varying neuronal activity in multiple connected brain areas, including the motor cortex (MCx), non-sensory thalamus, and others (Tanji and Evarts, 1976; Tanaka, 2007; Shenoy et al., 2013; Guo et al., 2017; Gao et al., 2018; Svoboda and Li, 2018). This ‘preparatory activity’ encodes specific upcoming movements, often seconds before movement onset (Tanji and Evarts, 1976). Cortical activity then changes rapidly and profoundly just before movement onset (Guo et al., 2014; Kaufman et al., 2016, 2014).

Recordings from large populations of neurons have enabled state-space analysis of neural activity in single trials. With *n* recorded neurons, the neuronal populations can be represented as a trajectory in *n*-dimensional activity space. These trajectories are often confined to a low-dimensional manifold, defined by several ‘activity modes’ that explain a significant proportion of the population activity. Activity modes are often obtained by projecting neural activity along specific directions in neural state space, or similar dimensionality-reduction methods (Cunningham and Yu, 2014; Kaufman et al., 2014; Kobak et al., 2016; Li et al., 2016). A successful decomposition provides activity modes that are interpretable, in the sense of predicting specific aspects of behavior (Mante et al., 2013; Kobak et al., 2016; Li et al., 2016; Inagaki et al., 2019; Finkelstein et al., 2019; Vyas et al., 2020; Lee and Sabatini, 2020).

For example, during motor planning, a preparatory activity in MCx occupies an activity mode that discriminates future movement types. This activity mode follows attractor dynamics and funnels preparatory activity to an initial condition (a fixed point) appropriate to trigger accurate and rapid movement (Churchland et al., 2010; Shenoy et al., 2013; Inagaki et al., 2019). After the Go cue, the dynamics in MCx shows large changes. The motor planning mode collapses (Funahashi et al., 1989; Shadlen and Newsome, 2001; Kaufman et al., 2016, 2014) and a new activity mode with multi-phasic dynamics emerges (Churchland et al., 2012). This movement type-specific mode is represented strongly in the descending MCx neurons that project to premotor neurons in the brainstem and spinal cord (Li et al., 2015; Economo et al., 2018) and presumably represents a motor command to initiate a specific movement. Another activity mode after the Go cue consists of changes that are invariant to the movement type (condition-invariant signal (Kaufman et al., 2016)), referred to here as ‘Go cue direction’ (GD) mode. Altogether, when an animal releases a planned action following a Go cue, neuronal activity in MCx transforms from a motor planning mode (i.e. preparatory activity) to a motor-command mode and a GD mode. These modes occupy different, near-orthogonal subspaces, which may explain in part why movements are not triggered during planning (Kaufman et al., 2014; Elsayed et al., 2016).

Neuronal dynamics underlying motor planning and execution have been studied in non-human primates and rodents trained in delayed-response tasks (Funahashi et al., 1989; Riehle and Requin, 1989; Erlich et al., 2011; Shenoy et al., 2013; Guo et al., 2014). An instruction informs movement type (e.g. movement direction or target; eye, tongue, arm or orienting movements) and a Go cue after a delay instructs movement onset: the Go cue releases planned actions. The anterior lateral motor cortex (ALM), a part of MCx, is necessary for motor planning and execution of directional licking in mice (Komiyama et al., 2010; Guo et al., 2014; Gao et al., 2018; Xu et al., 2019). Stimulation of ALM triggers rhythmic licking. ALM forms reciprocal connections with parts of the thalamus (referred to as ALM-projecting thalamus, or thal_ALM_) to maintain the motor plan (Guo et al., 2017). Thal_ALM_ receives input from the basal ganglia, cerebellum, and the midbrain, which directly or indirectly receive input from ALM (Guo et al., 2017; Gao et al., 2018). Thus, thal_ALM_ is a hub linking subcortical structures and ALM, forming multi-regional loops essential for orofacial movements.

In the context of a delayed directional licking task (Guo et al., 2014), we mapped the pathways and probed the mechanisms underlying cue-triggered switching of activity modes and the resulting movement initiation. By combining anatomy and large-scale, multi-regional electrophysiological recordings, we established the flow of Go cue-related information with millisecond time resolution from the midbrain to ALM via thal_ALM_. Ascending neurons in the pedunculopontine nucleus (PPN) and midbrain reticular nucleus (MRN) that project to thal_ALM_ show short-latency and transient GD signals. Optogenetic stimulation of these neurons parallels the effect of the Go cue, by initiating GD signals and motor command-like dynamics in ALM, followed by correct movements. Optogenetic silencing of these neurons abolished the GD signal and abolished behavioral responses. Altogether, we have identified a multi-regional pathway mediating cue-triggered mode switching that releases planned movement.

## Results

### A mode switch before movement initiation

We studied head-restrained mice performing a tactile delayed-response task (Guo et al., 2014) (Figures 1A and 1B, Movie S1). A tactile stimulus, an object presented to the right whiskers at one of two locations during the sample epoch, instructed lick direction (left or right). Mice were trained to withhold licking during the following delay epoch (1.2 s). After an auditory Go cue (3 or 3.4 kHz, 0.1s), licking in the correct direction was rewarded. In this task mice plan upcoming movements during the delay epoch and release planned movements following the Go cue.

**Figure 1.**
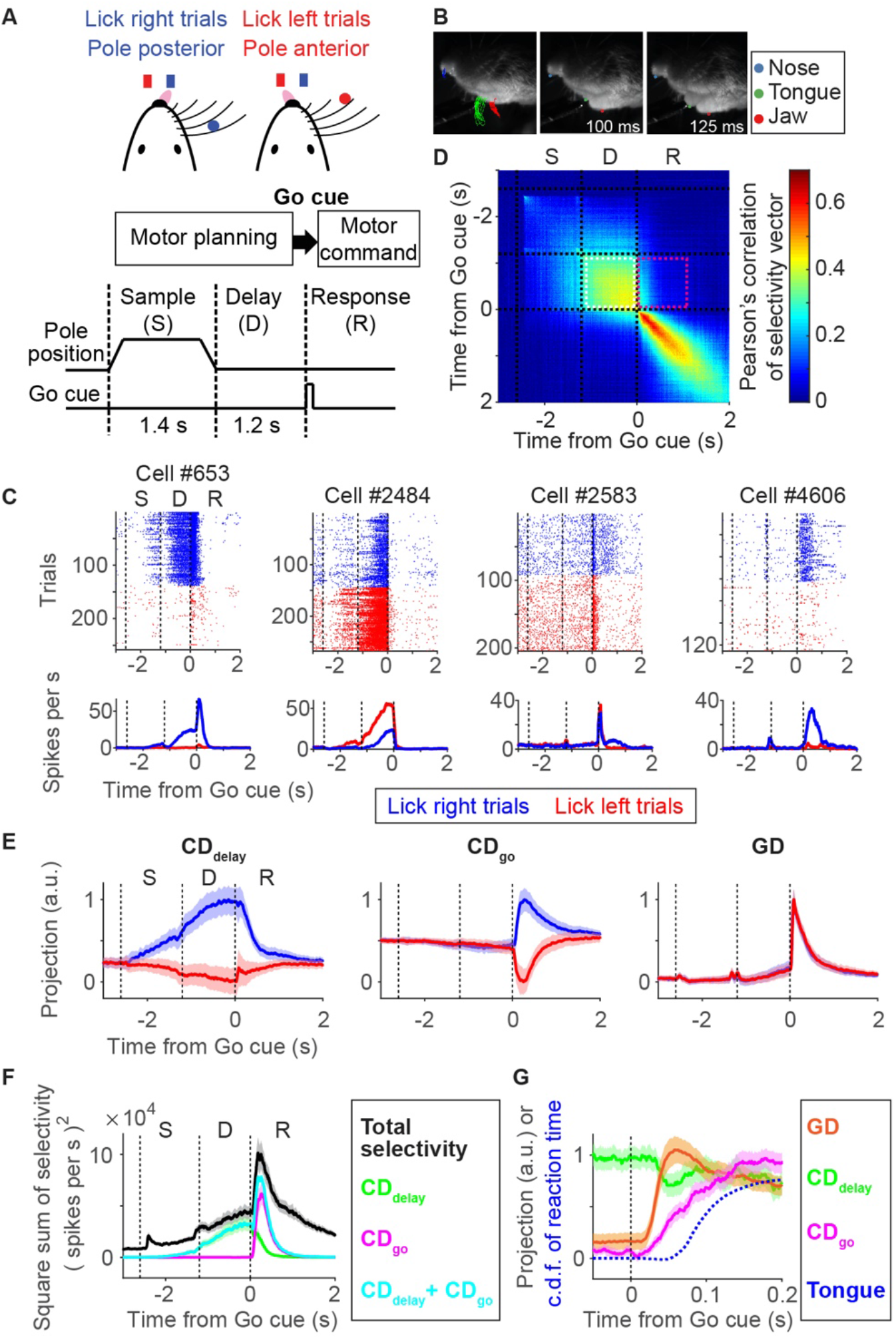
Activity modes for motor planning and movement initiation in anterior lateral motor cortex. A. Tactile delayed-response task. An object (pole, red / blue circle) is presented within reach of the whiskers during the sample epoch to instruct lick direction. B. Side view of a behaving mouse recorded with high-speed videography (400 Hz). Left, trajectories of nose (blue), tongue (green) and jaw (red) movement overlaid on an image at time 0 ms (onset of the Go cue). Middle, the first frame at which the tongue appears. C. Example neurons in ALM. Top, spike raster. Bottom, mean spike rate. Blue, correct lick right trials; red, correct lick left trials. Time is aligned to the timing of the Go cue. Dashed lines separate behavioral epochs. S, sample epoch; D, delay epoch; R, response epoch. D. Pearson’s Correlation of the population activity vector in ALM (bin: 10 ms; *n* = 5226 neurons). Dashed lines separate behavioral epochs. White dotted square, correlation between time points in the delay epoch; magenta dotted square, correlation between time points in the delay and the response epoch. E. Projections of activity along **CD**_**delay**_, **CD**_**go**_, and **GD**. Line, median. Shade, S.E.M. F. Selectivity explained by each direction in activity space. Black, square sum of selectivity of all recorded ALM neurons; Green, magenta, and cyan, the square sum of selectivity along **CD**_**delay**_, **CD**_**go**_, and their sum, respectively. G. Onset of each mode. Orange, activity along **GD** (mean of activity in lick right and left trials). Green and magenta, activity along **CD**_**delay**_ and **CD**_**go**_ (difference in activity between lick right and left trials). Dashed blue line, cumulative distribution of the first tongue detection (after the timing of the Go cue) by high-speed videography.

We performed extracellular recordings in left ALM (5923 neurons; we focused on 5226 putative pyramidal neurons for analyses). Consistent with previous reports (Guo et al., 2014; Inagaki et al., 2018), ALM neurons showed lick direction-selective spike rates (selectivity; *p* < 0.05, ranksum test) during the delay (2628/5226 neurons) and the response epochs (3271/5226 neurons). An early hypothesis suggested that preparatory activity is a subthreshold version of the activity that later causes the movement (Tanji and Evarts, 1976). This would imply that the Go cue enhances each neuron’s preparatory activity to trigger movement. Some neurons have activity consistent with this view. For example, cell #653 shows delay selectivity, and consistent selectivity peaks after the Go cue (Figure 1C). More generally, selectivity during the delay and response epoch was not consistent (Figures 1C, S1A, and S1D) (Kaufman et al., 2014). For example, cell #2484 shows lick-left selectivity during the delay epoch, but the selectivity collapses during the response epoch. Cell #4606 shows no selectivity during the delay epoch and strong lick-right selectivity during the response epoch (Figure 1C). These diverse neuronal activity patterns argue against the simple notion that preparatory activity is a subthreshold motor command.

To quantify how movement-related selectivity in ALM evolves at a population level we analyzed neural dynamics in activity space (Cunningham and Yu, 2014). We defined a population selectivity vector: ***w***_*t*_ = ***r***_lick-right, *t*_ − ***r***_lick-left, *t*_, where ***r***_lick-right, *t*_ and ***r***_lick-left, *t*_ are vectors of average spike rate of individual neurons for each time *t* in lick right and left trials, respectively (the length of the vector is the number of recorded neurons). Pearson’s correlation of this population selectivity vector is high across time points within the delay epoch (Figure 1D, a box with white dotted outline), implying that a similar combination of ALM neurons maintains selectivity during motor planning (Li et al., 2016; Economo et al., 2018; Inagaki et al., 2018). In contrast, population selectivity has a low correlation between time points before and after the Go cue (Figure 1D, magenta box), implying that different combinations of neurons show selectivity. Similarly, the population activity of ALM neurons within each trial type (***r***_lick-right, *t*_, or ***r***_lick-left, *t*_) shows low correlation before versus after the Go cue (Figure S1B). Moreover, intracellular recordings of ALM neurons show that membrane conductances increase rapidly after the Go cue, implying that neurons are clamped to a new state (Figures S1E and S1F). Altogether, these results imply that the population activity patterns in ALM change rapidly before and during movement initiation, similar to what has been observed across species and behavioral tasks (Funahashi et al., 1989; Shadlen and Newsome, 2001; Maimon and Assad, 2006; Vyas et al., 2020).

The stable preparatory activity during the delay epoch (Figures 1C, 1D, S1A and S1B) suggests a low-dimensional representation of ALM population activity. We defined a delay coding direction 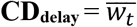 (−0.6 s < t < 0 s; time to Go cue) as the direction in activity space that discriminates future lick directions (lick left or right) during the delay epoch. Consistent with previous studies, this direction contains almost all movement direction-selective activity before the Go cue and allows decoding of lick direction one second before movement (Figures 1E and S1I) (Li et al., 2016; Inagaki et al., 2019). Similarly, we defined 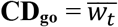 (0 s < t < 0.4 s; time to Go cue) as a direction that discriminates lick directions after the onset of the Go cue. We orthogonalized **CD**_**go**_ to **CD**_**delay**_ to characterize variance explained by each mode (Economo et al., 2018) (Figure S1G). Activity along the **CD**_**go**_ contains a large proportion of direction-selective activity and allows decoding of movement (Figures 1E and S1I). These two modes together explain 71.2 (65.3-76.0) % (mean, 2.5-97.5% confidence interval) of selectivity in ALM around the movement initiation (±200 ms from the Go cue; Figure 1F, cyan line).

Activities projected onto **CD**_**delay**_ and **CD**_**go**_ are correlated at the level of single trials (Figure S1K). This implies that information carried along **CD**_**delay**_ is transferred to **CD**_**go**_ following the Go cue (i.e., trials with strong activity along **CD**_**delay**_ have strong activity along **CD**_**go**_, and vice versa). This finding is consistent with the observation that fine scale movement parameters and reaction times are coded in preparatory activity (unpublished observations) (Li et al., 2016; Even-Chen et al., 2019) and shows that ALM preparatory activity (activity along **CD**_**delay**_) contributes to control of future movements (activity along **CD**_**go**_).

We also find phasic non-selective activity after the Go cue in ALM (e.g., cell #2583 in Figure 1C). At the population level we defined **GD** as the direction that discriminates activity before and after the Go cue (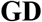 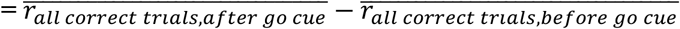; 100 ms time window). Activity along **GD** explains a large proportion of ALM activity after the Go cue (Figure S1H), similar to the condition-invariant signal found in a primate reaching task (Kaufman et al., 2016). Activity along **GD** is non-selective (Figure 1E) and cannot decode lick direction (Figure S1I), because activity changes around the Go cue are largely similar across trial types (Figure S1C). The trial-type differences that do exist contribute to **CD**_**go**_. These three directions in activity space, together with a fourth direction that captures non-selective ramping activity during the delay epoch (Li et al., 2016; Inagaki et al., 2018), account for 87.4 (84.8-89.6) % (mean, 2.5-97.5% confidence interval) of activity around the Go cue (±200 ms from the Go cue; Figure S1H, cyan line). Thus, these directions provide a near complete description of population activity around movement initiation.

Activities along **GD** and **CD**_**go**_ change rapidly after the Go cue (latencies, 20.0 (16-24) ms, 30.4 (18-44) ms, respectively; mean (2.5-97.5% confidence interval); Methods; Figure 1G) preceding movement onset (64.3 (56-75) ms; mean (2.5-97.5% confidence interval); blue dashed line in Figure 1G). Because activities along **GD** and **CD**_**go**_ precede movement (Figure S1J), and because silencing of ALM results in loss of behavioral responses (Komiyama et al., 2010; Gao et al., 2018; Xu et al., 2019), we hypothesized that activity along **CD**_**go**_ is the motor command as it is movement-type selective, whereas activity along **GD** switches the dynamics by terminating **CD**_**delay**_ and triggering **CD**_**go**_.

### CD_go_ is the motor command

To identify activity modes causal for movement execution, we analyzed ALM activity under conditions when mice failed to lick after the Go cue. First, we silenced ALM neurons projecting to the medulla (pyramidal tract neurons in lower Layer 5b, ‘PT_lower_’) (Economo et al., 2018) because the medulla contains the motor centers for orofacial movement (Travers et al., 1997; Stanek et al., 2014). To silence PT_lower_ cells, we injected AA_Vretro_ (Tervo et al., 2016) encoding soma-targeted (st) GtACR1 (Govorunova et al., 2015; Mahn et al., 2018) in the medulla (Figures 2A and S2A; Table S1 and S3). Bilateral optogenetic silencing of PT_lower_ cells in ALM (centered at AP 2.5 mm L 1.5 mm from Bregma; 1 second of photostimulation starting at the onset of Go cue) resulted in a loss of cue-triggered licking (Figures 2B, 2C, S2B and S2C). Similar bilateral silencing in posterior cortical regions (centered around AP 0 mm L 1.5 mm from Bregma) had a weaker behavioral effect (Figure S2C). Following the end of the photostimulus, mice licked in the correct direction (Figures 2B and S2D), implying that activity of ALM PT_lower_ cells is required to initiate movements, but not to maintain motor plans and not for a memory of the Go cue.

**Figure 2.**
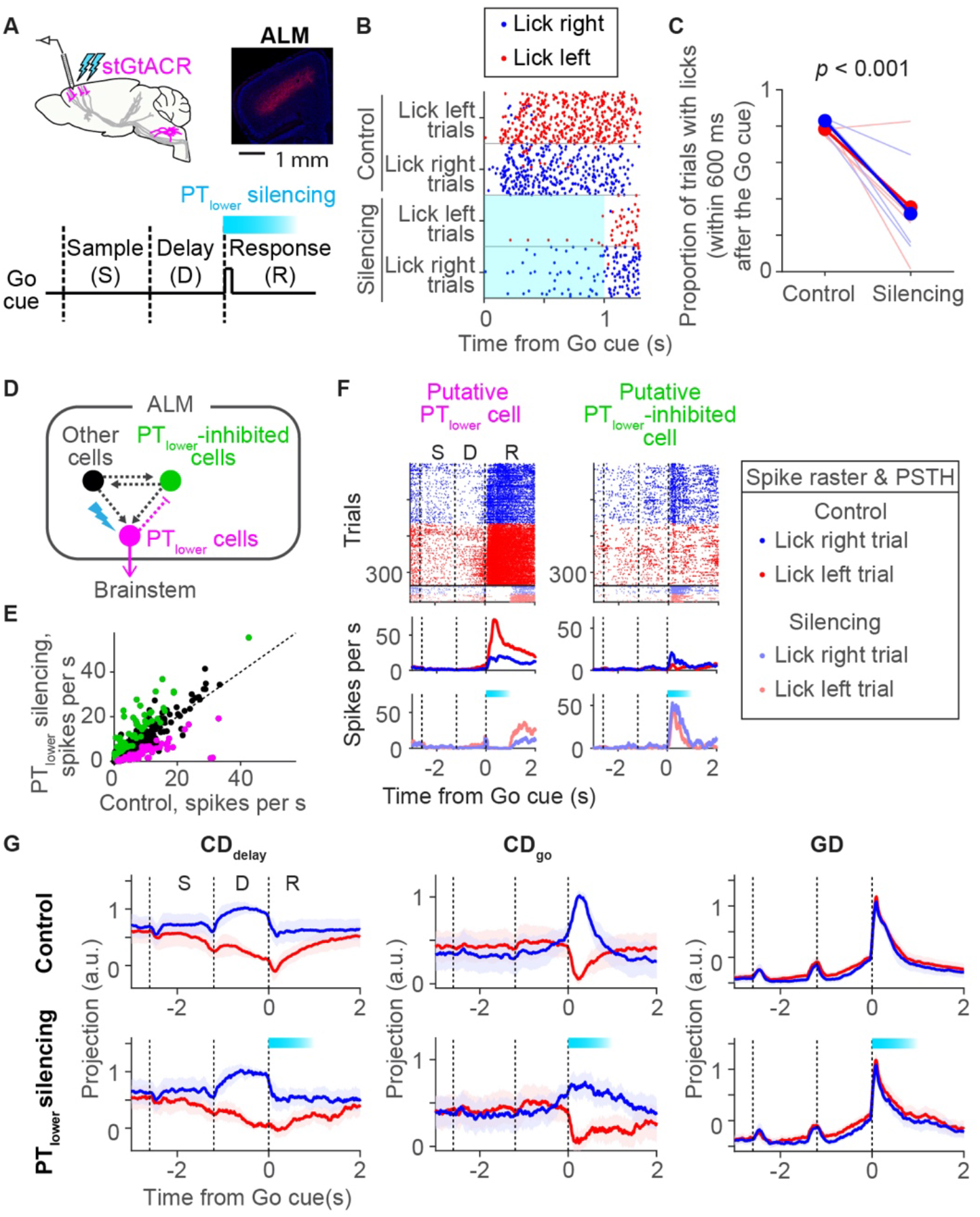
ALM motor commands are necessary for movement initiation. A. Silencing ALM neurons that project to orofacial motor centers in the medulla (PT_lower_). Inset, coronal section showing PT_lower_ neurons in ALM expressing GtACR1 fused with FusionRed; Blue, DAPI. B. Raster plot of lick timing in an example animal (75 trials shown per trial type). Cyan box, laser on. C. Proportion of licks within 600 ms after the Go cue. Blue, lick right trials; red, lick left trials. Circle, mean; lines, each animal, n = 4 mice. *P* < 0.001 in both lick right and left trials (hierarchical bootstrap with a null hypothesis that proportion of trials with licks in silencing trials are the same or higher than that in control). D. Schema showing PT_lower_ cells and other cell types (e.g. ‘PT_upper_’, brainstem-projecting neurons that do no innervate the medulla; intratelencephalic cells) (Li et al., 2015; Economo et al., 2018)analyzed in Figure 2 and S2. PT_lower_ cells (magenta) in the deep layers of ALM project to medulla. They indirectly inhibit PT_lower_-inhibited cells (green). E. Spike rate of individual neurons with or without PT_lower_ silencing. Circle, individual neuron; magenta; significantly decreased neurons (putative PT_lower_ cells); green, significantly increased neurons (putative PT_lower_ inhibited cells). F. Example putative PT_lower_ cell and putative PT_lower_-inhibited cell. Top, spike raster. Middle, PSTH in control trials. Bottom, PSTH in trials with PT_lower_ silencing. Blue, all lick right trials; red, all lick left trials: cyan bar, laser on. G. Projection of activity along **CD**_**delay**_, **CD**_**go**_, and **GD** with and without PT_lower_ silencing. Line, grand median across sessions (n = 24 sessions; 4 mice); shading, S.E.M. (*hierarchical bootstrap*); cyan bar, laser on.

We next performed extracellular recording in ALM during this manipulation, using a photostimulus with a significant behavioral effect (0.5 mW; Figure S2C). Spike rates were altered in 248/874 cells (*p* < 0.05, ranksum test; Figures 2D-2F, S2E and S2F). Cells silenced by the photostimulus (137/874) could include PT_lower_ cells and neurons coupled to them (Figures 2D-F; putative PT_lower_ cells), whereas excited cells (111/874) are neurons indirectly inhibited by PT_lower_ cells (Figures 2D-2F; putative PT_lower_-inhibited cells). Strongly silenced cells (*p* < 0.001, ranksum test) were in deep cortical layer (807 ± 151μm; mean ± std.; 51 cells), consistent with the depth distribution of PT_lower_ cells (Economo et al., 2018).

The PT_lower_ silencing attenuated activity along **CD**_**go**_ in lick-right trials (when recording in the left hemisphere), without affecting activity along other directions (**CD**_**delay**_ and **GD**) (Figures 2G and S2I). The contralateral reduction in **CD**_**go**_ does not simply reflect silencing of PT_lower_ neurons, but is a network effect. First, PT_lower_ cells are a small proportion of ALM neurons and thus make a correspondingly small contribution to **CD**_**go**_ (Figure S2A) (Economo et al., 2018). Second, putative PT_lower_ cells do not have a strong contralateral selectivity on average, and the extent of silencing is equal between trial types (i.e. no contralateral bias in putative PT_lower_ cells; Figures S2F and S2G). Third, **CD**_**go**_ based on non-PT_lower_ cells alone shows a contralateral reduction in **CD**_**go**_ activity during PT_lower_ silencing (Figure S2G). Although PT_lower_ cells have only weak connections with other pyramidal cells (Brown and Hestrin, 2009; Kiritani et al., 2012), they may influence the network via their connections to local GABAergic interneurons.

Additional support for **CD**_**go**_ in movement execution comes from analysis of trials in which mice failed to lick after the Go cue (no response trials; mostly near the end of a session when they are satiated). Activity along **CD**_**go**_ is attenuated without changes in other directions (during the first 200 ms after the Go cue; Figures S2H and S2I). Thus, activity along **CD**_**go**_ in ALM is necessary for movement initiation.

These experiments together show that descending output from ALM, PT_lower_ cells, is required for movement initiation. Furthermore, activity along **CD**_**go**_, which is encoded by PT_lower_ and other cells (Economo et al., 2018), precedes movement, instructs lick direction, and is required for movement execution. These results support a view that activity along **CD**_**go**_ is the motor command of ALM.

In contrast, activity along **GD** precedes movement but it is not instructive on movement type or sufficient to trigger movement by itself (without a change in activity along **CD**_**go**_). Thus, activity along **GD** is not a motor command. Instead, **GD** may trigger the activity along **CD**_**go**_ following the Go cue. To test this hypothesis, manipulations of activity along **GD** are necessary by activating or inhibiting neurons that carry this signal. This requires mapping the pathways that transmit the Go cue to ALM.

### Thalamus conveys the Go cue signal to ALM

To explore the causal chain of events leading from an auditory stimulus to movement initiation, we analyzed rapid changes in activity after the Go cue and compared latencies across brain areas. ALM forms strong reciprocal connections with thal_ALM_, including parts of the ventral medial (VM), ventral anterolateral (VAL), mediodorsal (MD), paracentral (PCN), central lateral (CL), central medial (CM), and parafascicular (PF) nuclei of the thalamus (Guo et al., 2017). The PCN, CL, CM, and PF comprise the so-called intralaminar (IL) nuclei of the thalamus.

We performed extracellular recordings in thal_ALM_ with high-density silicon probes and compared responses to the Go cue (i.e. changes in spike rate after the Go cue) to those in ALM (Figures 3A and 3B). Neurons in ALM and thal_ALM_ responded with increases or decreases in spike rate (go-up and go-down cells, respectively). The latency was shorter in thalamus (16.5 ± 1.5 ms; mean ± S.E.M.; time when 1% of cells show increase in spike rate) compared to ALM (25.0 ± 0.8 ms; mean ± S.E.M.) (*p* < 0.001; bootstrap; Figures 3A and 3B). The latency difference between thal_ALM_ and ALM is consistent with the action potential speed in thalamocortical projection neurons (Guo et al., 2017). Neurons with short latencies (< 20 ms) were wide-spread in thal_ALM_, and their spatial distribution was different from those of delay-selective neurons (Figure 3C) (Gaidica et al., 2018). We observed similar short latency activity in thal_ALM_ of mice performing an auditory delayed-response task (Figure S3) (Inagaki et al., 2018).

**Figure 3.**
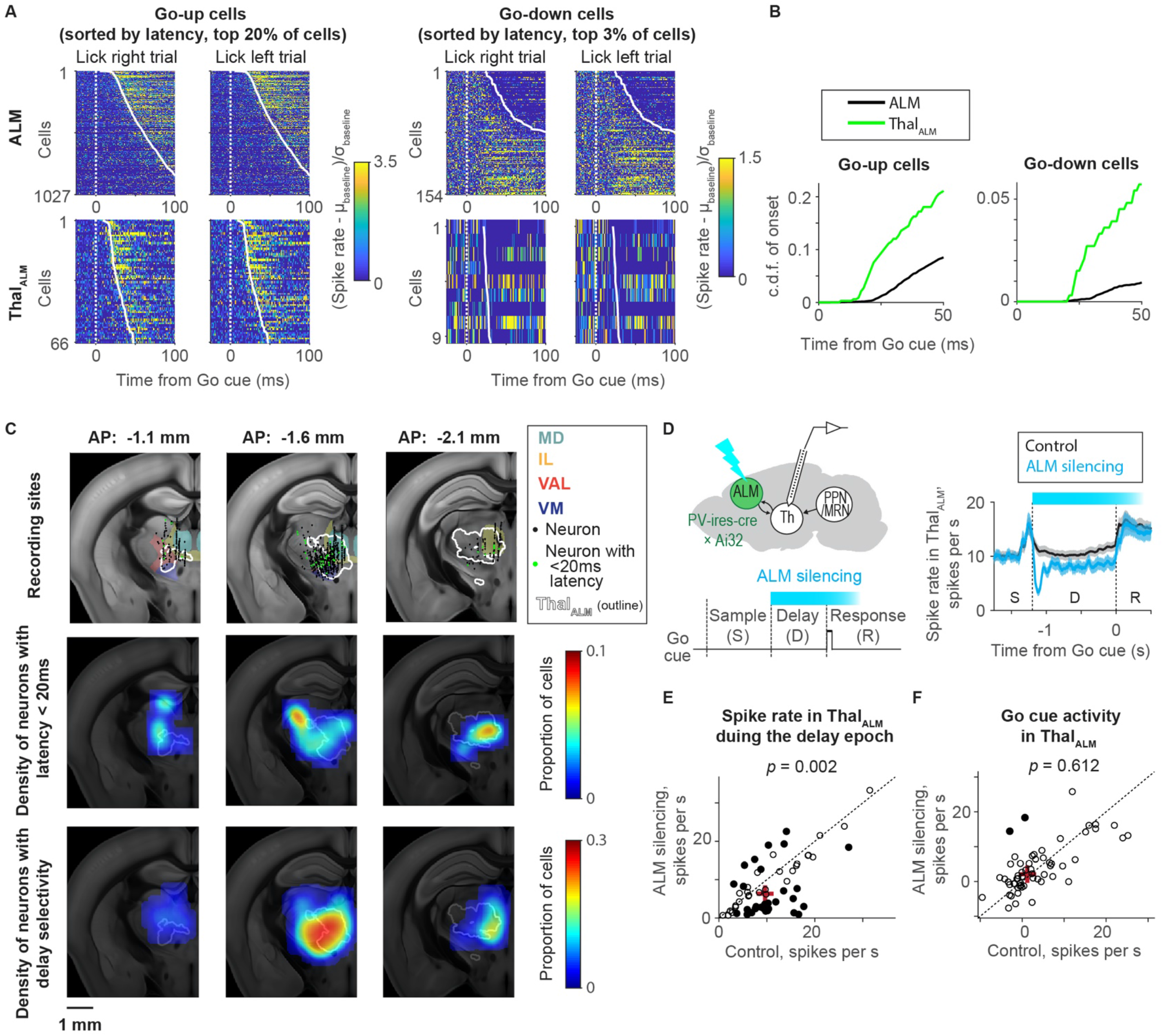
Short latency Go cue signals in ALM-projecting thalamus. A. Short-latency Go cue responses of ALM (top) and thal_ALM_ neurons (bottom) sorted by their latency. Neurons with increases (left, go-up cells) or decreases (right, go-down cells) in spike rate are shown separately. The fastest 20% and 3% cells are shown for go-up and go-down cells, respectively. Spike rates are normalized by the baseline (spike rate before the Go cue, 100 ms window) for each neuron. μ, mean; *σ*, standard deviation. B. Cumulative distribution function (c.d.f.) of latency to the Go cue across neurons in ALM and thal_ALM_. C. Recordings in thalamus. Top, recording sites in Allen Common Coordinate Framework (CCF). Each colored region corresponds to a different thalamic nucleus. White contour, thal_ALM_. Black dots, location of individual recorded neurons. Green, neurons with < 20 ms latency. Middle, the density of neurons with latencies < 20 ms. Bottom, the density of neurons with delay selectivity. AP, posterior to Bregma. D. Recording in thal_ALM_ during ALM silencing. Left, schema. Right, mean activity of thal_ALM_ with or without ALM silencing. Cyan bar, photoinhibition of ALM. E. Spike rate during the delay epoch in thal_ALM_ with or without ALM silencing. Circle, individual neuron in thal_ALM_ (n = 58 cells). Filled circle, significantly modulated cells (*p* < 0.01, ranksum test). Cross, median activity. *P-value*, signrank test comparing spike rate across neurons with or without silencing. F. The amplitude of Go cue activity (spikes per s; change in spike rate after the go cue; 100 ms window) in thal_ALM_ with or without ALM photoinhibition. The same format as in **E**.

Photoinhibition of ALM reduced the activity of thal_ALM_ during the delay epoch (Guo et al., 2017) (Figures 3D, 3E, and S3E). It did not change the amplitude of the Go cue response in thal_ALM_, although the photoinhibition lasts until the response epoch (Figure 3F). Thus, ALM is not necessary for the Go cue response in thal_ALM_. Together with the latency analysis, these results indicate that the Go cue activity first arrives in thal_ALM_, then drives ALM (Dacre et al., 2019).

Auditory cortex neurons respond to sounds with short latencies (12 ms; (Williamson and Polley, 2019)) but do not directly project to ALM (Guo et al., 2017; Oh et al., 2014). M1 (AP 0.15 mm, ML 1.7 mm from Bregma) and ALM are bidirectionally connected (Guo et al., 2017). M1 showed latencies similar to ALM (Figure S3; 20.6± 5.1 ms; mean ± S.E.M.; *p* = 0.41, Bootstrap). Because M1 is not necessary for initiation of directional licking (Xu et al., 2019), and because of slow inter-cortical interactions between M1 and ALM (Guo et al., 2017), a parsimonious explanation is that the Go cue response in ALM does not rely on M1. Altogether, thal_ALM_ signals the Go cue to ALM, yet additional contributions from M1 cannot be excluded.

### Inputs to ALM-projecting thalamus

To identify inputs to thal_ALM_, we injected retrograde tracers in thal_ALM_ (retrobeads and AAV_retro_, Figures S4A-S4C). Both tracers labeled cells in the ipsilateral frontal cortex and multiple subcortical areas such as substantia nigra pars reticulata (SNr), superior colliculus (SC), deep cerebellar nuclei (DCN, composed of dentate nucleus, DN; interposed nucleus; and fastigial nucleus, FN), and PPN/MRN (Figures 4A and S4A-S4C), consistent with previous reports (Saper and Loewy, 1982; Krout et al., 2002; Martinez-Gonzalez et al., 2011; Guo et al., 2017; Gao et al., 2018).

**Figure 4.**
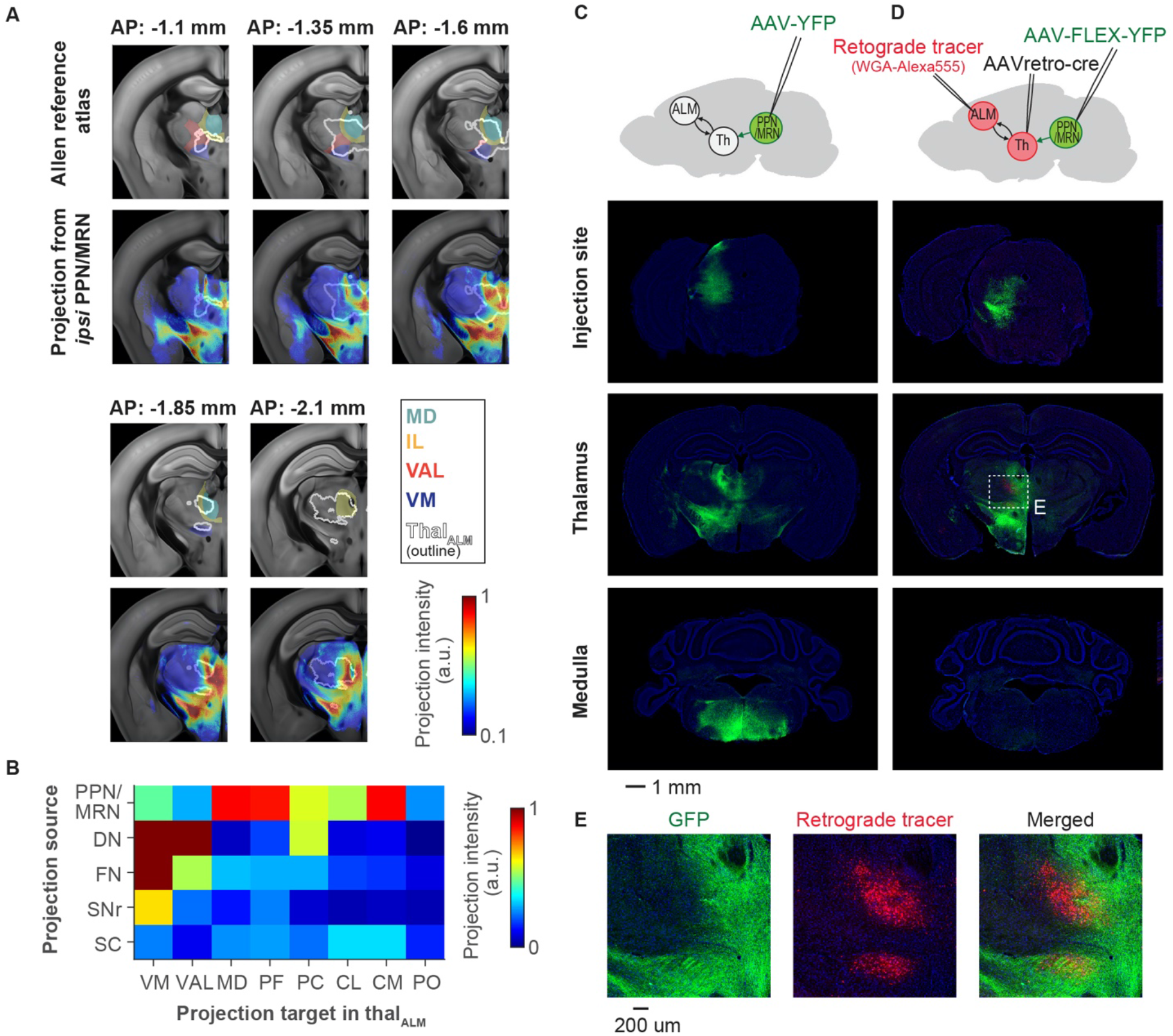
PPN/MRN projects to ALM-projecting thalamus. A. Projections from ipsilateral PPN/MRN to thal_ALM_ (coronal view). Top, Allen Reference Atlas. Each colored region corresponds to a different thalamic nucleus. White contour, thal_ALM_. Bottom, the intensity of projection from PPN/MRN, registered to the Allen CCF (mean of 4 mice). AP, posterior to Bregma. B. Quantification of anterograde labeling in thal_ALM_ from subcortical structures (Methods). C. Top, projection-type non-specific labeling of PPN/MRN neurons. Bottom, signal at the injection site, in the thalamus, and in the medulla. The image gains and contrasts are identical between the images of the thalamus and the medulla. Green, YFP; Blue, Nissl staining. D. Top, labeling thalamus-projecting PPN/MRN neurons. Bottom, same as in **C** for thalamus-projecting PPN/MRN neurons. Red, retrograde tracing from ALM. E. Enlarged image of the thalamus in **D**.

To map the projections from these subcortical areas within thal_ALM_ and beyond we performed anterograde labeling by injecting AAVs expressing fluorescent proteins in each area (Figures 4A and S4D; Table S2). PPN/MRN neurons have wide-spread projections to thal_ALM_, whereas other structures have relatively localized projections (Figure 4B). We focused on PPN/MRN because the output extensively overlaps with the wide-spread short latency Go cue responses in thal_ALM_ (Figure S4F).

Thalamus-projecting PPN/MRN (PPN/MRN_Th_) neurons are distributed across a region referred to as the ‘mesencephalic locomotor region’ (MLR) (Shik et al., 1966). Stimulation of the MLR produces gait changes in cats and rodents, which is mediated by glutamatergic neurons that descend into the medulla (Shik et al., 1966; Roseberry et al., 2016; Josset et al., 2018; Caggiano et al., 2018). PPN/MRN neurons in mice project to both thalamus and the medulla (Figure 4C). To label PPN/MRN_Th_ specifically, we injected AAV_retro_-Cre in thal_ALM_, and Cre-dependent YFP-expressing AAV in PPN/MRN (Figure 4D). By injecting a retrograde tracer in ALM of the same animal, we confirmed that projections from PPN/MRN partially overlap with thal_ALM_ (Figure 4E). These experiments revealed that PPN/MRN_Th_ neurons lack a projection to the medulla, and thus constitute a distinct intermingled population from neurons that project to the medulla (Figure 4D). PPN contains a high density of cholinergic (*chat*+) cells, and a subset are known to project to thalamus (Sofroniew et al., 1985; Mena-Segovia and Bolam, 2017; Huerta-Ocampo et al., 2020). However, the majority (75%) of PPN/MRN_Th_ neurons are glutamatergic (*vglut2*+), not Chat+ or GABAergic (*gad1*+), as confirmed by fluorescent *in situ* hybridization and immunohistochemical staining (Figures S4G-S4J; Table S4). We used whole-cell patch-clamp recordings in acute brain slices to confirm direct glutamatergic input from PPN/MRN to thal_ALM_ neurons (Figures S4K-S4P).

### Latency after the Go cue in thalamus-projecting brain areas

Next, we compared the latencies of Go cue responses across the subcortical areas projecting to thal_ALM_. We analyzed extracellular recordings in DCN (Gao et al., 2018) and SNr (Guo et al., 2017), and performed extracellular recordings centered around SC and PPN/MRN, all in the same task (Figure 5A).

**Figure 5.**
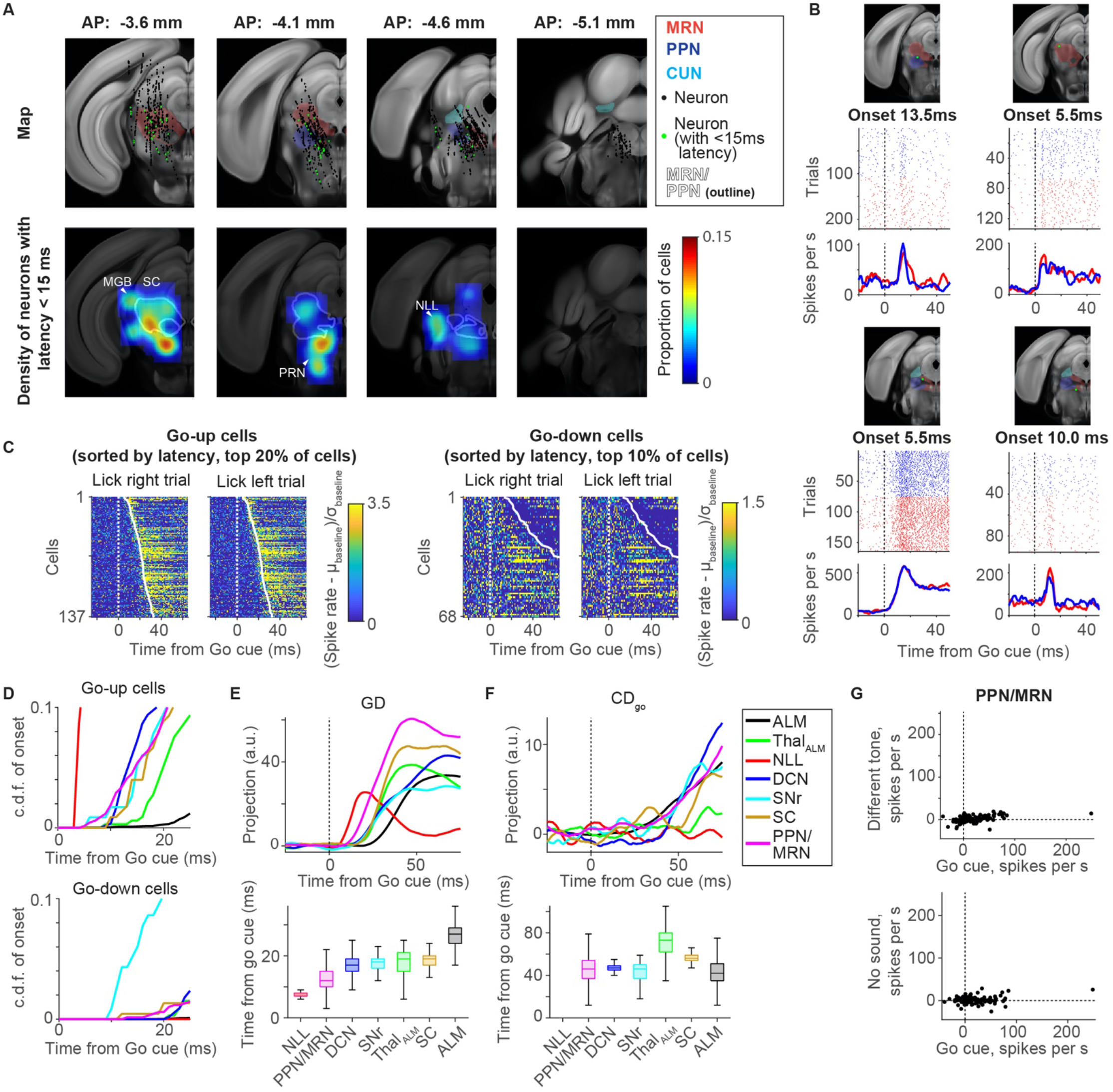
Short latency Go cue signals in PPN/MRN. A. Recordings in the midbrain. Top, recording sites in Allen CCF. Each colored region corresponds to a different midbrain nucleus. Black dots, location of individual recorded neurons. Green, location of neuron with <15 ms latency. Bottom, the density of neurons with latency to the Go cue shorter than 15 ms. White contour indicates MRN and PPN. CUN, cuneiform nucleus. PRN, pontine reticular nucleus. B. Four example PPN/MRN neurons. Top, location of recorded neuron (green circle) in the Allen CCF. Middle, spike raster. Bottom, mean spike rate. C. Go cue response of PPN/MRN sorted by their latency to the Go cue. Neurons with an increase or decrease in spike rate after the Go cue are shown separately. Spike rates are normalized by the baseline (spike rate before the Go cue, 100 ms window) for each neuron. μ, mean; *σ*, standard deviation. D. Cumulative distribution (c.d.f.) of latency to the Go cue across neurons in each brain area (see Methods for number of neurons analyzed). Each color indicates a different brain area (box next to **F**). E. Top, projection of activity along **GD** in each brain area. Bottom, the latency of post-Go cue increases in activity along **GD** (Methods). Central line in the box plot is the median. Top and bottom edges are the 75% and 25% points, respectively. The whiskers show the lowest datum within 1.5 interquartile range (IQR) of the lower quartile, and the highest datum within 1.5 IQR of the upper quartile. F. Same as **E** for activity along **CD**_**go**_. G. Increase in spike rate of PPN/MRN neurons in response to the Go cue and different tone (top) or no sound at the expected timing of the Go cue (bottom). Circles, neurons (n = 178).

We found neurons with fast Go cue responses (<15 ms) in multiple areas. This includes the nucleus of lateral lemniscus (NLL), an auditory center that receives direct input from the cochlear nucleus (Davis et al., 1982); the pontine reticular nucleus, which is known to be part of the acoustic startle pathway (Davis et al., 1982); the auditory thalamus (medial geniculate body, MGB) (Figure 5A; arrowheads). In addition, we found sparsely distributed cells with short latency Go cue responses in PPN/MRN (white dashed outline; Figures 5A-5C) (Reese et al., 1995; Dormont et al., 1998; Pan et al., 2005).

We compared the latencies and spike rates after the Go cue across brain areas (Figures 5D and S5). All analyzed subcortical areas contain cells with Go cue latencies shorter than those in thal_ALM_. To analyze the latency of the non-selective component of the Go cue response, we projected activity in each area to its **GD**. Among the subcortical areas projecting to thal_ALM_, activity along **GD** first emerged in PPN/MRN after the Go cue on average (Figure 5E). Unlike activity along **GD**, the selective component of the Go cue response, i.e. activity along **CD**_**go**_ in each brain area, emerged almost simultaneously across brain areas, and later than **GD** (Figures 5F). Thus, following the Go cue, non-selective activity rapidly spreads across brain areas, followed by emergence of selective activity.

The fast Go cue responses in PPN/MRN are not simple auditory responses. Rather, they constitute a learned contextual response that is specific to the sound used as the Go cue. We tested this by recording in mice trained with either 3 or 12kHz Go cue and trained to ignore the other tone (Figures 5G and S5G). The response does not reflect timing of the task because there was no response without the Go cue (Figure 5G). Go cue responses in thal_ALM_ were also specific to the Go cue, consistent with a view that PPN/MRN signals to thal_ALM_ (Figure S5G).

Although all thalamus-projecting brain areas contained cells with Go cue latencies shorter than thal_ALM_, since PPN/MRN contains the cells with the shortest latencies (Figures 5D and 5E), and since their projections overlap with a subregion of thal_ALM_ with the fast Go cue responses (Figure S4F), we next tested whether activity in thalamus-projecting PPN/MRN is causal for the cue-triggered activity in ALM and movement initiation.

### Phasic stimulation of thalamus-projecting PPN/MRN neurons mimics the Go cue

If PPN/MRN signals the Go cue to ALM via thalamus, phasic optogenetic stimulation of the PPN/MRN_Th_ should trigger the effects of the Go cue. To stimulate PPN/MRN_Th_ specifically, we injected ChR2-expressing AAV in PPN/MRN (unilaterally or bilaterally, n =20 mice) and placed a fiber optic unilaterally in thal_ALM_ for axonal stimulation (Figures 6A and S6A-S6C) (Petreanu et al., 2007). Mimicking the phasic Go cue response with brief (5 or 10 ms) photostimulation of PPN/MRN axons increased licking responses (Figures 6B and S6H; *p* < 0.001, bootstrap; differences across animals are partially explained by the location of AAV injection in PPN/MRN and by the evoked activity patterns in ALM; Figures S6D-S6F). When mice licked in response to the stimulation, they licked in the correct direction defined by the trial type (Figure 6C). This was the case even in mice with unilateral injection of ChR2 expressing AAV in PPN/MRN and ipsilateral photostimulation in thal_ALM_ (Figure S6G; n = 3 mice; note that PPN/MRN neurons project to ipsilateral Th, and more weakly to contralateral thalamus; Figure S4E). Releasing the movement to the direction instructed by the tactile cues is precisely what the Go cue does: it does not carry directional information by itself, but releases planned movements. This property of PPN/MRN stimulation is unusual, as unilateral stimulation of motor-related areas generally drives unidirectional movements. For example, stimulation of MCx, basal ganglia, motor thalamus, SC triggers contralateral movement (Li et al., 2015; Dacre et al., 2019; Lee et al., 2020; Lee and Sabatini, 2020), whereas stimulation of DCN triggers ipsilateral movement (Gao et al., 2018).

**Figure 6.**
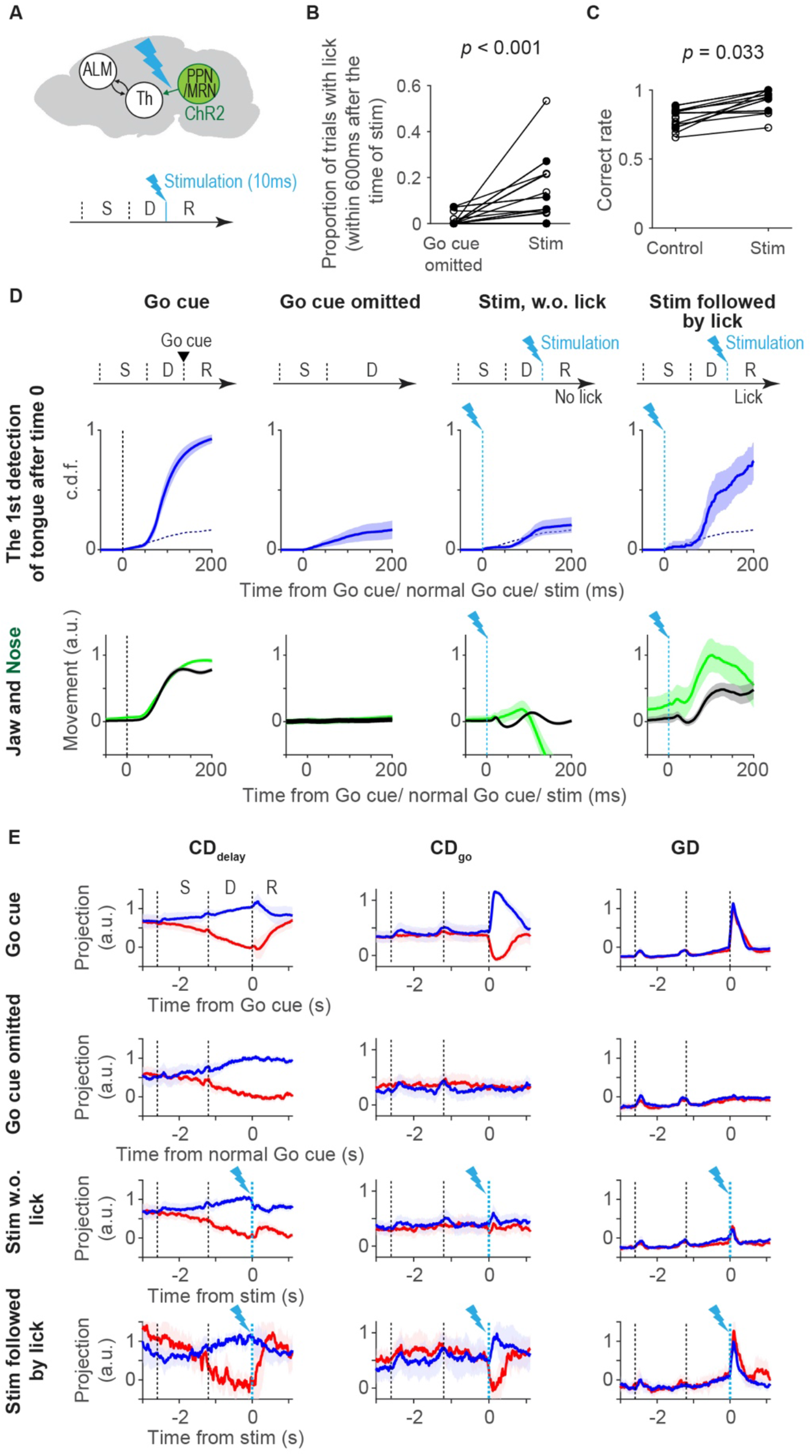
Stimulation of thalamus-projecting PPN/MRN neurons triggers planned movement. A. Schema of PPN/MRN_th_ stimulation experiment. B. Proportion of trials with lick after PPN/MRN_th_ stimulation. Circle, mouse (n = 20 mice). Filled circle, mice with unilateral virus injection (n = 4 mice). *P-value*, hierarchical bootstrap with a null hypothesis that the proportion of trials with licks in stimulation trials are the same or lower than that in control. C. Same as **B** for the proportion of trials with correct lick direction (correct rate) after the Go cue (control) or stimulation. D. Top, cumulative distribution of the first tongue detection (with high-speed videography) after time 0. Dotted line, data in Go cue omitted condition for comparison. Bottom, jaw movement (black), and nose movement (green). Trials are classified as follows: Go cue, trials with the Go cue; Go cue omitted, trials without the Go cue or stim; Stim w.o. lick, trials with stimulation but without lick; Stim followed by lick, trials with stimulation followed by lick. Tongue detection onset: (Go cue) 64.3 (56.0-75.0) ms; mean (2.5-97.5% confidence interval); (stim followed by lick) 75.5 (50.0-118.0) ms; *p* = 0.194 (hierarchical bootstrap). Jaw movement onset: (Go cue) 33.2 (20.0-42.5) ms; (stim followed by lick) E. 69.7 (8.8-102.5) ms; *p* = 0.114 (hierarchical bootstrap). Nose movement onset: (Go cue) 43.0 (32.5-50.0) ms; (stim followed by lick) 61.9 (7.5-117.1) ms; *p* = 0.139 (hierarchical bootstrap). The null hypothesis for *p*-value is that the onset of Go cue trials is shorter than that in stim trials. F. Projection of activity along **CD**_**delay**_ (left), **CD**_**go**_ (middle), and **GD** (right) across trial types. Cyan dashed line, photo-stimulation. Line, grand median of sessions (n = 21 sessions; 12 mice); shading, S.E.M. (*hierarchical bootstrap*).

In addition, photostimulation of PPN/MRN_Th_ induced orofacial movements similar to those triggered by the Go cue (monitored by high-speed videography; Figures 1B and 6D; Movie S2). Mice did not initiate movement by anticipating the Go cue timing (Figure 6D; Go cue omitted). In trials in which mice did not lick in response to the stimulation, mice showed attenuated orofacial movements (Figure 6D; stim, w.o. lick).

We next compared ALM activity with Go cue-triggered licks and PPN/MRN_Th_ stimulation-triggered licks. We performed extracellular recordings in ALM. The changes in activity produced by the Go cue and photostimulation were remarkably similar. In particular, the photostimuli did not merely excite all ALM neurons by increasing glutamatergic input from PPN/MRN_Th_. Instead, in trials in which mice licked in response to photostimulation, neural activity in ALM resembled activity triggered by the Go cue at the level of individual cells (Figure S6I) and at the population level (Figures S6J and S6K) by both increasing and decreasing spike rates (Dacre et al., 2019).

To analyze the population activity pattern further, we projected the population activity along **CD**_**delay**_, **CD**_**go**_, and **GD**. The activity underlying stimulation-triggered licks was similar to that underlying cue-triggered licks. Namely, we observed a significant change in activity along **GD** and **CD**_**go**_ (Figures 6E and S6L). Furthermore, activity along **CD**_**delay**_ collapsed after the photostimulation, similar to changes observed after the Go cue (Figure 6E). These changes happen before the onset of movement (within 50 ms after the onset of stimulation; Figures S6L and S6M), implying that they are not caused by the movement. In trials in which mice did not lick in response to photostimulation, changes in activity were much attenuated, consistent with the reduced orofacial movements (Figures 6D and 6E). No activity change was observed in Go cue omitted trials consistent with the lack of movement.

The amplitude of activity along **GD** predicted whether photostimulation triggered licks on a trial-by-trial basis (Figure S6N; *p* < 0.05 in 4 out of 7 sessions). Activity along **CD**_**go**_ is determined by both activity along **GD** and **CD**_**delay**_ (Figures S6O and S6P). Altogether, when photostimuli induced sufficiently large activity along **GD** they create activity along **CD**_**go**_ in a manner proportional to the activity along **CD**_**delay**_, which results in licking in the planned direction.

### Optogenetic perturbation of thalamus-projecting PPN/MRN neurons blocks movement initiation

To test whether activity in PPN/MRN_Th_ neurons is necessary for cue-triggered movement initiation, we performed perturbation experiments. We expressed stGtACR1 in PPN/MRN_Th_ bilaterally by injecting AAV_retro_-hsyn-Cre in thalamus and AAV-FLEX-stGtACR1 in PPN/MRN (Figures 7A, S7A and S7B). Optogenetic perturbation during the Go cue presentation blocked cue-triggered licks (Figure 7B; laser was on 0.6 s before the Go cue and lasted for 1.2 s; 473 nm 40 Hz sinusoidal). Perturbation of CamKII+ PPN/MRN_Th_ (using AAV_retro_-CamKII-Cre) resulted in a similar behavioral effect, implying that CamKII+ PPN/MRN_Th_ cells are necessary for the cue-triggered movement initiation (Figure 7B). *In situ* hybridization confirmed that CamKII+ cells are glutamatergic (Figure S4H) (Roseberry et al., 2016). In contrast, perturbation of cholinergic neurons, a common cell type in PPN, did not affect movement initiation (Figure 7B). Altogether, glutamatergic PPN/MRN_Th_ neurons are necessary for movement initiation. Even after the end of the perturbation, mice showed reduced licks (Figure S7C). Thus, unlike PT_lower_ silencing in which mice licked after silencing (Figure 2B), mice behaved as if there was no Go cue.

**Figure 7.**
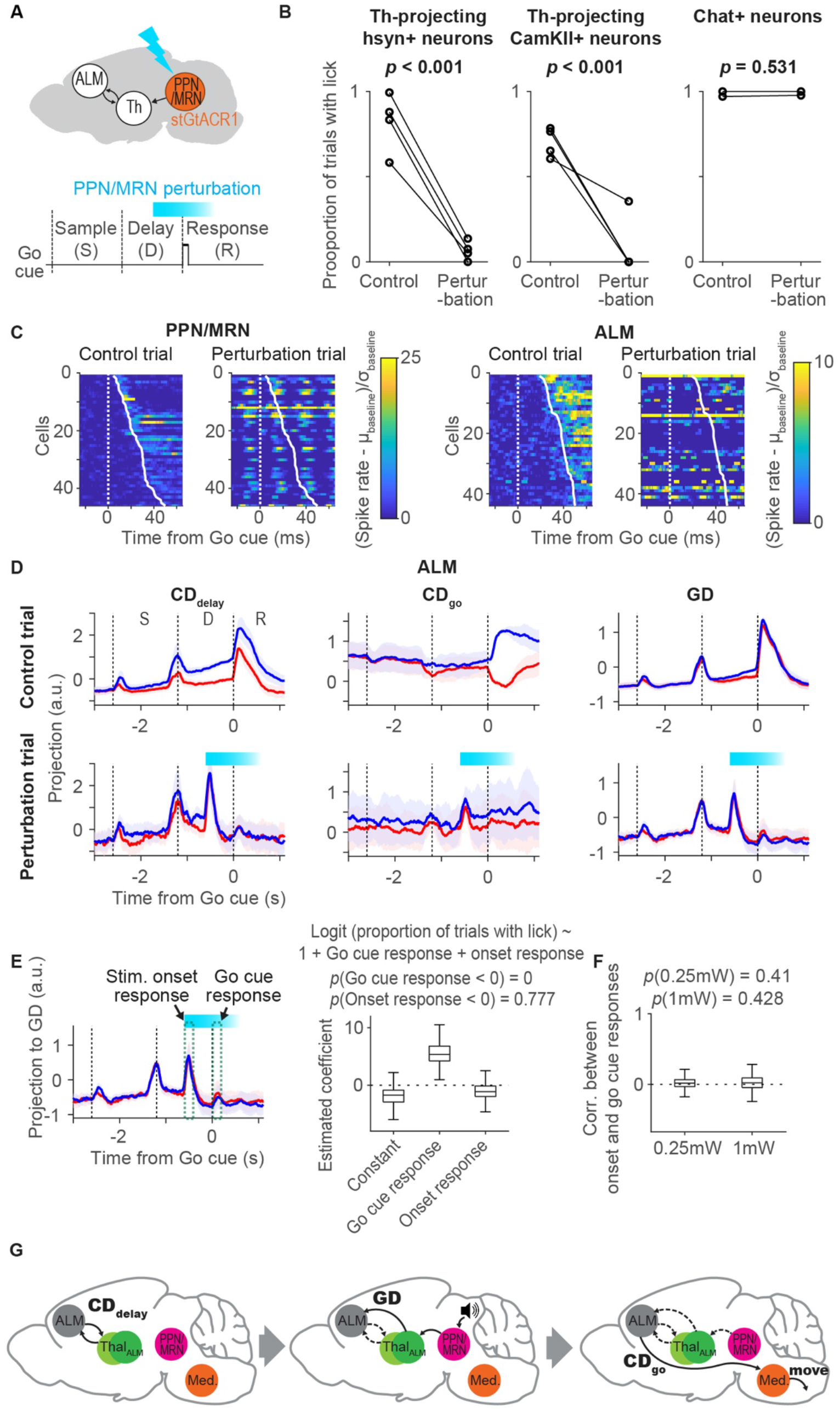
Activity in thalamus-projecting PPN/MRN neurons is required for cue-triggered movement initiation. A. Perturbation of PPN/MRN_th_ before and during the Go cue. B. Behavioral effects of perturbing Th-projecting hsyn+ neurons (left; n = 4 mice), Th-projecting CamKII+ neurons (middle; n = 4 mice), and Chat+ neurons (right; n = 2 mice; note Chat+ cells are not necessarily projecting to thalamus) in PPN/MRN. *P-value*, hierarchical bootstrap with a null hypothesis that the proportion of trials with licks in perturbation trials are the same or higher than that in control. C. Go cue response sorted by their latency to the Go cue. Neurons with increase in spike rate before movement (within 50 ms after the Go cue) are shown (45/292 cells in PPN/MRN and 44/635 cells in ALM). Activities of the same neurons in control (left) and perturbation (right) trials. Spike rates are normalized by baseline (spike rate before the Go cue in control trials, 100 ms window). In **C**-**F**, data of perturbing Th-projecting CamKII+ neurons is shown. D. Projection of activity along **CD**_**delay**_, **CD**_**go**_, and **GD** across trial types. Cyan, laser on. Line, grand median of sessions (n = 17 sessions; 4 mice); shading, S.E.M. (*hierarchical bootstrap*). E. Trial-by-trial relationship between proportion of trials with lick, activity along **GD** at the Go cue (go cue response) and at the stim onset (onset response). Left, schema of activity analyzed in the regression analysis. Mean activity within the green dotted lines were analyzed (window size, 200 ms). Right, estimated coefficients of logit regression. *P-value*, hierarchical bootstrap (n = 17 sessions; 4 mice). F. Correlation between activity along **GD** at the Go cue and at the stim onset. *P-value*, hierarchical bootstrap with a null-hypothesis that coefficient is lower than 0 (n = 17 sessions; 4 mice). G. Multi-regional flow of information underlying the cue-triggered movement initiation. Left, preparatory activity (**CD**_**delay**_) is maintained in a cortico-thalamocortical loop. Middle, the Go cue (speaker) activates PPN/MRN, which then activates neurons in thal_ALM_, which are different from neurons that maintain preparatory activity (green circles). This induces activity along **GD** in ALM. Right, the **GD** activity then causes a collapse of activity along **CD**_**delay**_ and an emergence of motor command (**CD**_**go**_), which engages medulla (Med.) circuits to initiate planned movements.

To confirm the effect of the optogenetic manipulation, we performed extracellular recordings in mice expressing stGtACR1 in CamKII+ PPN/MRNTh. A subset of PPN/MRN neurons showed increases in spike rate time-locked to the sinusoidal laser modulation (23/45 cells; *p*<0.05, ranksum test; analyzed cells with latency shorter than 50 ms), presumably due to axonal excitation caused by stGtACR1 (Figures 7C and S7D) (Mahn et al., 2018; Messier et al., 2018). Indeed, increases in spike rate occurred around and slightly after the peak of the laser power (Figure S7F). Other PPN/MRN neurons (13/45; *p* < 0.05, ranksum test) were silenced (Figures 7C and S7D). In both cases, the Go cue response was abolished in perturbation trials (Figures 7C and S7D).

Next, we performed extracellular recordings in ALM. Like in PPN/MRN, the Go cue response was lost in individual ALM neurons during the perturbation (Figures 7C and S7E). The sinusoidal modulation was attenuated in ALM (Figure S7F). ALM neurons showed a loss of Go cue response when hsyn+ or CamKII+ PPN/MRN_Th_ neurons were perturbed, but not when Chat+ PPN/MRN were perturbed, consistent with the behavioral effect (Figure S7G).

In the activity space, the Go cue response disappeared across all directions (Figures 7D and S7I). In addition to the loss of the Go cue response, the photostimulus onset caused transient activity (onset response), which was likely induced by stGtACR1-dependent axonal excitation (Figure 7E) (Mahn et al., 2018; Messier et al., 2018). We used logistic regression to confirm that the loss of licking is not caused by the onset response. This analysis shows that larger attenuation of the Go cue response corresponds to lower probability of licking (Figure 7E). In addition, the amplitude of the onset response was not correlated with the loss of Go cue response (Figures 7F and S7H). Altogether, a parsimonious explanation is that the perturbation of the Go cue response in PPN/MRN results in a loss of Go cue response in ALM and a loss of movement initiation.

## Discussion

Activity in the anterior lateral motor cortex (ALM) is critical for motor planning and execution of directional licking in mice (Komiyama et al., 2010; Guo et al., 2014; Gao et al., 2018; Xu et al., 2019). During a delayed-response task, ALM activity can be decomposed into a several activity modes that capture a large proportion of cortical activity (Figure 1) (Li et al., 2016; Inagaki et al., 2018). During the delay epoch, movement-selective preparatory activity is contained mostly in the **CD**_**delay**_ direction in activity space. After the Go cue, this activity rapidly reorganizes into the non-selective **GD** mode and the direction-selective **CD**_**go**_ mode. This progression is necessary to initiate movement (Figure 2).

We identified a multi-regional neural pathway that is critical for reorganizing ALM activity in response to the Go cue and to initiate planned directional licking (Figure 7G). Ascending glutamatergic neurons in PPN/MRN signal Go cue information to ALM via thal_ALM_ and thereby cause the reorganization of ALM activity and release planned movements. Our conclusions are based on multiple lines of evidence. First, PPN/MRN contains neurons that specifically respond to the auditory Go cue (and not to other sounds) (Figure 5G). Second, latencies after the Go cue are shorter in PPN/MRN than in thal_ALM_ (Figure 5E). Third, PPN/MRN neurons project to areas of thal_ALM_ that show short-latency Go cue responses (Figure 4). Fourth, brief optogenetic stimulation of PPN/MRN_Th_ neurons triggers rapid changes in ALM activity, similar to those caused by the Go cue itself (Figure 6E). Fifth, optogenetic stimulation of PPN/MRN_Th_ neurons elicited the appropriate directional licking, even after unilateral stimulation of PPN/MRN_Th_ neurons (Figure 6C), unlike other thal_ALM_ projecting areas. Sixth, perturbation of PPN/MRN_Th_ activity resulted in a loss of Go cue response in ALM and abolished behavioral responses (Figure 7). Our findings are consistent with previous experiments in other species. For example, in rats and monkeys, bilateral lesion of PPN/MRN blocks cue-triggered movement initiation (Wilson, 1973; Florio et al., 1999). In humans, PPN/MRN and downstream thalamic areas show activity in cued movement tasks (Kinomura et al., 1996).

PPN/MRN receives projection from NLL (Reese et al., 1995), which itself receives direct auditory input from the cochlear nucleus (Davis et al., 1982). Consistent with this view, the latency to the Go cue in NLL is shorter than in PPN/MRN (Figure 5). The auditory response in PPN/MRN is specific to the sound associated as the Go cue (Figure 5G). PPN/MRN is likely the site of this association.

Although we emphasize the PPN/MRN_Th_ → thal_ALM_ → ALM circuit here, PPN/MRN neurons likely exert their role in movement initiation via additional brain regions. For example, PPN/MRN_Th_ neurons also project to the SNr and subthalamic nucleus (STN) (Figure 4A) (Martinez-Gonzalez et al., 2011; Vitale et al., 2019). STN in turn projects to SNr, and SNr is known to control premotor circuits in the superior colliculus and medulla (DeLong, 1990; Hikosaka et al., 2000; Klaus et al., 2019). In addition, it is possible that PPN/MRN_Th_ locally excite or inhibit PPN/MRN neurons that descend to motor centers. Indeed, PPN/MRN is ideally positioned to coherently control brain-wide circuits in cortex, basal ganglia, and the brainstem for movement initiation. Our experiments also do not exclude contributions from additional subcortical areas to movement initiation, such as DCN (Spidalieri et al., 1983; Gemba and Sasaki, 1987; Gao et al., 2018; Dacre et al., 2019) and basal ganglia (da Silva et al., 2018; Díaz-Hernández et al., 2018).

Most of our understanding of thalamocortical processing is based on sensory thalamus and cortex. Much less is known about non-sensory (‘higher-order’) thalamus and its interactions with the frontal cortex. Cortex and higher-order thalamus are strongly coupled in both directions (Bolkan et al., 2017; Guo et al., 2017; Schmitt et al., 2017). Our experiments suggest that midbrain sends simple contextual signals to the cortex via particular thalamic nuclei to modulate cortical activity modes. Different thalamic nuclei contain neurons with different projection patterns (Steriade et al., 1997; Clascá et al., 2012). For example, VM contains neurons that have broad projection patterns to layer 1 (‘matrix’). VAL instead projects in a more focal manner to middle layers (‘core’) (Jones, 1998; Kuramoto et al., 2015). The ‘intralaminar’ nuclei project to deep cortical layers and the striatum. These different thalamocortical projections activate cortical microcircuits in specific ways (Anastasiades et al., 2020). We find that the spatial distribution of thal_ALM_ neurons with short latencies to the Go cue differs from those showing delay selectivity (Figure 3C). It will be interesting to learn how the thalamic nuclei showing different activity patterns modulate cortex through their specific thalamocortical projections. We note that the PPN/MRN input to thal_ALM_ produces short-latency and reliable synaptic currents with paired-pulse depression (Figure S4P). Thus, the PPN/MRN_Th_ → thal_ALM_ projection has the hallmarks of a classic driver input (Guillery and Sherman, 2002).

Animal behavior often consists of multiple phases, each corresponding to different computations. Activity modes underlying each phase likely occupy near-orthogonal activity subspaces so that the different computations do not interfere with each other (Mante et al., 2013; Kaufman et al., 2014; Stavisky et al., 2017; Hennig et al., 2018; Rouse and Schieber, 2018). However, the information carried by a mode needs to be transferred to subsequent modes to mediate coherent behavior. For example, information encoded by the preparatory activity must be transferred to the motor command to initiate planned movement. Indeed, we observed correlation between activity along **CD**_**delay**_ during the delay epoch and **CD**_**go**_ after the Go cue (Figures S1K and S6P).

We designed our optogenetic manipulations to test the causal roles of each activity mode. PT_lower_ neurons project to motor centers that control orofacial movements in the medulla (Li et al., 2015; Economo et al., 2018). Silencing PT_lower_ neurons resulted in a loss of movements and reduced activity along **CD**_**go**_, without affecting **GD** (Figure 2G). Optogenetic stimulation of PPN/MRN_Th_ axons first increased activity along the **GD**. When **GD** activity was sufficiently large it also induced selective activity along **CD**_**go**_ and triggered appropriate behavioral responses (Figures 6E and S6O). Thus, although both **CD**_**go**_ and **GD** emerge after the Go cue, they are dissociable: activity along **GD** is not sufficient to trigger movement by itself. Instead, it induces activity along **CD**_**go**_, which then controls the movement via PT_lower_ cells.

Our analysis relies on the millisecond temporal precision provided by electrophysiology and behavioral tracking with high-speed video, as well as a behavioral task with multiple choices. Our results provide a clear demonstration that state space analysis can extract features of population activity that have specific roles in behavior.

Parkinson’s disease (PD) patients experience difficulty in self-initiating movement, clinically described as freezing of gait (FOG). However, they can often perform complex movements in response to sensory cues, such as catching a ball. This phenomenon, known as paradoxical kinesis, is commonly used for rehabilitation (Ginis et al., 2018). Neurodegeneration in PD impacts activity in basal ganglia, a structure important for motor control (DeLong, 1990; Gerfen and Surmeier, 2011; Hikosaka et al., 2000; Klaus et al., 2019; Mink, 1996). PPN/MRN_Th_ → thal_ALM_ → MCx (e.g. ALM) pathway may initiate movement bypassing basal ganglia (Schwab et al., 2020), which could explain why cue-triggered movement is spared in PD patients. In addition, deep brain stimulation (DBS) of PPN has been applied to treat the FOG in PD (Thevathasan and Moro, 2019). The PPN DBS improves simple reaction tasks to sensory cues (Thevathasan and Moro, 2019), raising a possibility that PPN DBS is acting on the cue-triggered movement initiation mechanism (i.e., by affecting PPN/MRNTh). Further investigation on genetically-defined cell types and precise location of PPN/MRN neurons that underlie cue-triggered movement may help to optimize treatment of PD.

## Acknowledgements

We thank Svoboda and Inagaki lab members and B. Sauerbrei for comments on the manuscript, M. N. Economo. and S. Romani for helpful discussions, M. Inagaki for animal training, T. Harris, B. Barbarits, J. Colonell, B. Karsh, W. L. Sun, J. J. James, A. Liddell, and M. Pachitariu for help with silicon probe recordings and spike sorting, T. Wang for help with HCR, L. Narayan for help with high-speed videography, K. Ritola for viruses, and M. Mahn and O. Yizhar for GtACR plasmids. This work was funded by ZIA MH002487-30 (C.R.G.), NOW-Vidi and ERC-Stg grants (Z.G.), Australian National Health and Medical Research Council and the Australian Research Council: CE140100007 (P.S.), Searle Scholars Program (N.L. and H.K.I.), NIH NS112312 (N.L.), Simons Collaboration on the Global Brain (K.S. and N.L.), Wellcome Trust (S.C.), Helen Hay Whitney Foundation (H.K.I.), Max Planck Florida Institute (H.K.I.) and Howard Hughes Medical Institute (K.S.).

## Author Contributions

H.K.I. and K.S. planned the study. H.K.I. and S.C. performed experiments. H.K.I. analyzed data with S.C., K.S. and Z.Y. M.R. and P.S. performed acute slice recordings. C.G., H.H., and Z.G. performed histology. N.L. provided DCN recordings. H.K.I. and K.S. wrote the paper, with input from all the authors.

## Declaration of interests

The authors declare no competing interests.

## Methods

### Mice

This study is based on both male and female mice (age > P60 except for acute slice recording). We used eight mouse lines: C57Bl/6J (JAX #000664), VGAT-ChR2-EYFP (JAX #14548) (Zhao et al., 2011), PV-IRES-Cre (JAX #017320) (Hippenmeyer et al., 2005), Chat-IRES-Cre (JAX #006410) (Rossi et al., 2011), Vglut2-IRES-Cre mice (JAX #028863) (Vong et al., 2011), Sst-IRES-Cre (JAX #013044) (Taniguchi et al., 2011), Rbp4-Cre (MMRRC031125, Gerfen et al., 2013), and Ai32 (JAX #024109) (Madisen et al., 2012). See Supplementary Table 1 and 2 for mice used in each experiment.

All procedures were in accordance with protocols approved by the Janelia Institutional Animal Care and Use Committee or the University of Queensland Animal Ethics Committee. Detailed information on water restriction, surgical procedures, and behavior have been published (Guo et al., 2014a). Mice were housed in a 12:12 reverse light: dark cycle and behaviorally tested during the dark phase. A typical behavioral session lasted 1 to 2 hours. Mice obtained all of their water in the behavior apparatus (approximately 1 ml per day; 0.3 ml was supplemented if mice drank less than 0.5 ml). Mice were implanted with a titanium headpost. For cortical photoinhibition, mice were implanted with a clear skull cap (Guo et al., 2014b). Craniotomies for recording were made after behavioral training.

### Virus and tracer injection

We followed (dx.doi.org/10.17504/protocols.io.bctxiwpn) for virus and tracer injection. See Supplementary Table 1 and 2 for descriptions of viruses and injection coordinates used for each experiment. We used the following tracers: WGA-Alexa555 (WGA-Alexa Fluor® 555; Thermo Fisher Scientific) and Red RetroBeads (Lumafluor). See Supplementary Table 3 for a list of viruses used in this research.

### Behavior

For the tactile delayed-response task (Guo et al., 2014a) (all experiments but Figure S3A-S3D), at the beginning of each trial, a metal pole (diameter, 0.9 mm) moved within reach of the whiskers (0.2 s travel time) for 1.0 s, after which it was retracted (0.2 s retraction time). The sample epoch (1.4 s total) was the time from the onset of pole movement to the completion of pole retraction. The delay epoch lasted for another 1.2 s after the completion of pole retraction. An auditory ‘Go cue’ separated the delay and the response epochs (pure tone, 3 or 3.4 kHz, 0.1 s).

A two-spout lickport (4.5 mm between spouts) was used to record licking events and deliver water rewards. After the Go cue, licking the correct lickport produced a water reward (approximately 2 μL); licking the incorrect lickport triggered a timeout (0 to 5 s). Licking before the Go cue (‘early lick’ trials) was punished by a timeout (1 s). Trials in which mice did not lick within 1.5 seconds after the ‘go’ cue (‘no response’ trials) were rare and typically occurred at the end of behavioral sessions.

For Figure 5G, individual mice were trained with 3 (or 12) kHz (pure tone, 0.1 s) Go cues. In addition, they were trained to ignore tones with different frequency, 12 (or 3) kHz, played at the time of normal Go cue (different tone trials). Go cue omitted trials (trials without Go cue) and different tone trials were deployed in 25% randomly selected trials. Licking in Go cue omitted trials and different tone trials was punished with a timeout (1 s).

For the auditory delayed-response task (Figure S3A-S3D), tones were presented at one of two frequencies: 3 or 12 kHz, during the sample epoch. Each tone was played three times for 150 ms with 100 ms inter-tone intervals. The following delay epoch lasted for another 1.2 seconds. An auditory ‘go’ cue (carrier frequency 6 kHz, with 360 Hz modulating frequency, 0.1 s) separated the delay and the response epochs.

### Optogenetics

Photostimulation was randomly deployed on ~25% behavioral trials. To prevent mice from distinguishing photostimulation trials from control trials using visual cues, a ‘masking flash’ (1 ms pulses at 10 Hz) was delivered using 470 nm LEDs (Luxeon Star) throughout the trial. For both ChR2 and stGtACR1, we used a 473 nm laser (Laser Quantum). The laser power was controlled by an acousto-optical modulator (AOM; Quanta Tech) and a shutter (Vincent Associates). See Supplementary Table 1.

The ChR-assisted photoinihibition of ALM (Figures 3D-3F) was performed through clear-skull cap (beam diameter at the skull: 400 μm at 4 σ) (Guo et al., 2014b). We stimulated parvalbumin-positive interneurons in PV-IRES-Cre mice crossed to Ai32 reporter mice expressing ChR2 (Guo et al., 2017) for 1.6 s starting at the onset of the delay epoch (T_delay_) with 200 ms ramping down (mean laser power: 1.5mW). We silenced ALM ipsilateral to the recorded thalamus.

To silence medulla-projecting ALM neurons (PT_lower_) bilaterally (Figure 2), we photoinhibited for 1 s with 100 ms ramping down, starting at the timing of the Go cue. We photoinhibited four spots in each hemisphere, centered on ALM (AP 2.5 mm; ML 1.5 mm) with 1 mm spacing (in total eight spots bilaterally) using scanning Galvo mirrors through clear-skull cap. We photoinhibited each spot sequentially, at the rate of 5 ms per spot (period, 20 ms). The laser powers noted in the figures and text indicate the mean laser power per spot.

For PPN/MRN_Th_ axonal photostimulation experiments (Figure 6), we randomly interleaved three trial types: (1) Go cue trials, trials with Go cue at T_delay_ + 1.2 s (i.e. 1.2 s after delay onset; this is the control condition mice were trained with, which constitutes 75~85% of trials during the experiments); (2) Go cue omitted trials, trials without Go cue; (3) stimulation trials, trials with axonal excitation of Thalamus-projecting PPN/MRN by 20mW 473nm laser at T_delay_ + 1.2 s for 5 or 10 ms (through N.A. 0.37 fiber optics; see Supplementary Table 1 for coordinates). In both Go cue omitted trials and stimulation trials, a delayed Go cue was presented at T_delay_ + 2.4 s and licks to this Go cue were rewarded in order to maintain behavioral performance.

To prevent mice from associating optogenetic stimulation with water reward (and increasing licks because of such association), we did not provide water reward to stimulation-triggered licks (licks after the stimulation and before the delayed Go cue). Consequently, mice decreased stimulation-triggered licks over trials/sessions, presumably by learning to distinguish stimulation and the actual Go cue (Figure S6H).

For PPN/MRN_Th_ perturbations using stGtACR1 (Figure 7), we delivered photostimuli bilaterally to PPN/MRN_Th_ starting at T_delay_ + 0.6s lasting 1.2s duration (with 200 ms ramp up and down to minimize axonal excitation, 40Hz sinusoidal modulation; Go cue was presented at T_delay_ + 1.2 s). We tested 0.25, 1, and 10 mW photostimuli. The strongest photostimulus triggered strong axonal excitation, and was excluded from the analyses.

Note that stGtACR1-driven synchronous excitation of PPN/MRN_Th_ (Figures S7D and S7F) could have potentially propagated to the thalamus to shunt other subcortical input. If the thalamic activity were entrained by PPN/MRN_Th_ input, we would expect to see similar activity patterns in ALM because of the tight thalamocortical coupling (Guo et al., 2017). However, we did not see this (Figures S7E and S7F). In addition, the behavioral effect of PPN/MRN_Th_ perturbation was explained by the stGtACR1-mediated loss of the Go cue response in ALM but not by stGtACR1-driven excitation (Figure 7). Thus, a parsimonious explanation is that PPN/MRN_Th_ perturbation suppressed the Go cue signals, which then resulted in the loss of Go cue response in ALM and behavior.

### Behavioral data analysis

To calculate the proportion of trials with licks, ‘lick early’ trials were excluded (Figures 2C, 7B, S2C, and S2D). To calculate the correct rate (i.e., the proportion of licks to the correct direction), ‘lick early’ trials and ‘no response’ trials were excluded (Figures 6C, S2C, and S6G). ‘Lick’ was defined as a contact of the tongue with the electrical lick ports. This is why there are trials with tongue movement (based on high-speed videography) without lick (Figure 6D, Stim w.o. lick).

For the PPN/MRN stimulation experiment (Figure 6B), we plotted the proportion of trials with T_delay_ + 1.2 s < T_lick_ < T_delay_ + 1.8 s (T_lick_ denotes the timing of the first lick). Since stimulation was delivered at T_delay_ + 1.2s, this corresponds to a proportion of trials with licks within 0.6 s after the stimulation. An increase in lick during stim trials (Figures S6D, S6F, and S6H) is the difference in the proportion of trials with licks between stim trials and the Go cue omitted trials. To calculate the correct rate of stim trials (Figures 6C and S6G, right), we considered lick directions of licks within T_delay_ + 1.2 s < T_lick_ < T_delay_ + 1.8s. The control (Figures 6C and S6G, left) is the correct rate of Go cue trials.

For statistics, we performed hierarchical bootstrapping: first, we randomly selected animals with replacement, second, randomly selected sessions of each animal with replacement, and third randomly selected trials within each session with replacement.

### Videography and analysis

High-speed videography of orofacial movement (side view) was taken at 400 Hz using a CMOS camera (acA2040-180km, Basler) with IR illumination (940nm LED, Roithner Laser). We used DeepLabCut (Mathis et al., 2018) to track the movement of the tongue, jaw, and nose (Figure 1B; Supplementary movies 1 and 2). For the jaw and nose, movements along dorsoventral direction were analyzed. The amplitude of movement was normalized per session so that the mean position at T_delay_ + 1.2 seconds (time 0 in Figure 6D) was 0 and the maximum movement in the Go cue trials was 1. Note that the jaw moves downward after the Go cue, but due to this normalization, the value increases in Figure 6.

To calculate the onset of jaw and nose movement, we performed hierarchical bootstrapping: first, we randomly selected animals with replacement; second, randomly selected sessions of each animal with replacement; third, randomly selected trials within each session with replacement. Then, we calculated the mean trace of these randomly selected trials. Next, we linearly detrended the mean trace based on its activity between T_delay_ + 0.6 s and T_delay_ + 1.2 s (time 0 to −0.6 in Figure 6D). We identified the time point in which movement exceeds three times the standard deviation of the baseline before the Go cue (100 ms window). We repeated this procedure 1000 times to estimate the mean and S.E.M.

To calculate the onset of tongue movements, first, we calculated the cumulative distribution (c.d.f.) of the first time point tongue was detected by DeepLabCut after the Go cue (T_delay_ + 1.2 seconds). We subtracted the c.d.f. of a trial type of interest by the c.d.f. of the Go cue omitted trial (Figure 6D, dotted line). The movement onset is the time point at which the difference passes 0.05. We repeated this procedure with hierarchical bootstrapping 1000 times to estimate the mean and S.E.M.

### In vivo whole-cell recording and analysis

All recordings were made from the left ALM. Data and detailed procedures have been published (Guo et al., 2017). In brief, we partially compensated for series resistance and injected a ramping current until action potentials disappeared (Anderson et al., 2000; Yu et al., 2016) (767 ± 172 pA for positive current injection, −164 ± 64 pA for negative current injection; mean ± standard deviation). To calculate mean membrane potential, spikes were clipped off. Only correct trials were analyzed.

### Extracellular electrophysiology

A small craniotomy (diameter, 0.5 – 1 mm) was made over the recording sites one the day prior to the first recording session. Extracellular spikes were recorded using Janelia silicon probes (HH-2) with two shanks (32 channels each, 25 μm interval between channel, 250 μm between shanks), or Neuropixels probes (Jun et al., 2017b; Steinmetz et al., 2020; Liu et al., 2020). For the HH-2 probe, 64 channel voltage signals were multiplexed, recorded on a PCI6133 board (National instrument), and digitized at 400 kHz (14 bit). The signals were demultiplexed into 64 voltage traces sampled at 25 kHz and stored for offline analysis. All recordings were made with open-source software SpikeGLX (http://billkarsh.github.io/SpikeGLX/). During recording, craniotomy was immersed in cortex buffer (NaCl 125 mM, KCl 5 mM, Glucose 10 mM, HEPES 10 mM, CaCl_2_ 2 mM, MgSO_4_ 2 mM, pH 7.4).

Brain tissue was allowed to settle for at least five minutes before recording started. For ALM, recording depth (between 800 μm to 1100 μm) was inferred from manipulator readings. For subcortical areas, electrode tracks labelled with CM-DiI were used to determine recording locations (Liu et al., 2020) (Figure S5H).

### Extracellular recording data analysis

JRClust (Jun et al., 2017a) with manual curation (all data except the recordings during the auditory delayed-response task; Figure S3A-S3D) or Kilosort2 (https://github.com/MouseLand/Kilosort2) were used for spike sorting.

For the tactile delayed-response task, we recorded 9472 neurons across 300 behavioral sessions from 53 mice. For ALM, in total, 6030 neurons were recorded across 173 behavioral sessions from 37 mice. For the thalamus, in total, 640 neurons were recorded across 23 behavioral sessions from 4 mice. For the midbrain, 2808 neurons were recorded across 102 sessions from 18 mice. In addition, we analyzed published data collected from mice trained with the same tactile delayed-response task: 611 thalamus neurons and 116 SNr neurons from (Guo et al., 2017), and 554 DCN neurons from (Gao et al., 2018). Spike widths were computed as the trough-to-peak interval of the mean spike waveform. In ALM, putative pyramidal neurons (units with spike width > 0.5 ms) were analyzed (Guo et al., 2014b). In thalamus and SNr, units with width > 0.35ms and < 0.35ms, respectively, were analyzed following the criteria in (Guo et al., 2017).

For the auditory delayed-response task (Figure S3A-S3D), we obtained 13139 units across 76 sessions from 15 animals. We analyzed 5072, 655, 607, 1145, and 1560 units in ALM, M1, thal_ALM_, SC, and PPN/MRN, respectively. Spike sorting results of Kilosort2 were not curated manually. Instead, we used a combination of quality metrics (https://github.com/AllenInstitute/ecephys_spike_sorting) to extract potential good units for analysis. Specifically, we selected units with amplitude > 100 μV, ISI violation < 0.5, amplitude cutoff < 0.1, SNR > 2.5, spike width < 1.2 ms, and a presence ratio > 0.95 over the course of recording sessions.

For Figures 1, 3, and 5, neurons with at least 40 correct lick-right trials and 40 correct lick-left trials were analyzed. Forty trials were randomly subsampled for each correct trial type (lick-right and left) to analyze latency and selectivity. To calculate the latency from the Go cue for each neuron, we assumed that spike generation follows a *Poisson* process. First, we calculated the baseline spike rate before T_go_ (time of the Go cue; 100 ms window to calculate the baseline). Then, identify the first time point after T_go_ in which spike rate becomes higher or lower than a significance level, α = 0.001 (*Poisson* distribution based on the baseline spike rate; we call this time point as T_*p* = 0.001_). We defined the latency as the last time-point in which the spike rate becomes higher or lower than a second significance level, α = 0.05 (*Poisson* distribution) between (T_go_, T_*p*=0.001_]. We took this two steps approach to avoid detecting small amplitude changes.

Delay- and response-selective cells (Figure S5C) are neurons with significant delay or response selectivity (ranksum test comparing spike counts in correct lick right vs. correct lick left trials during (T_go_ – 0.6s, T_go_), and (T_go_, T_go_ + 0.6s), respectively; *p* < 0.05.)

For the peri-stimulus time histograms (PSTHs) in Figures 1C, 5B, and S1A, only correct trials were included. In Figures 2F, S2E, S6I, S7D, and S7E, correct, incorrect, and no-lick trials were pooled to compare control vs. perturbation conditions. PSTHs were smoothed by a 100 ms boxcar filter for plots including all epochs (e.g., Figure 1B) or by a 5 ms causal boxcar filter for plots zoomed in around the Go cue (e.g., Figure 5B). In Figures 3A, 5C, 7C, and S5A, spike rates were standardized by the mean and standard deviation of spike rates before the Go cue (100 ms window).

To plot the estimated density of neurons in Figures 3C and 5A, we applied a 2D Gaussian filter (standard deviation = 250 μm) to the estimated location of each neuron. Then, for each pixel, we calculated the density as (number of neurons with short latency or delay selectivity) / (number of neurons + 0.05). To prevent pixels with low numbers of neurons from having an extremely high estimated density, we added 0.05 to the denominator.

### Coding direction analysis

To calculate delay coding direction (**CD**_**delay**_) for a population of *n* recorded neurons, we looked for an n × 1 vector that maximally distinguished the two trial types in the *n-dimensional* activity space. For each time point *t*, we defined a population selectivity vector: ***w***_*t*_ = ***r***_lick-right, *t*_ − ***r***_lick-left, *t*_, where ***r***_lick-right, *t*_ and ***r***_lick-left, *t*_ are n × 1vectors of average spike rate of individual neurons in correct lick right and left trials without optogenetic manipulations (unperturbed correct trials). We averaged *wt* during the last 600 ms of the delay epoch (T_go_ - 0.6 s < t < T_go_) and normalized by its own norm to obtain the **CD**_**delay**_. To calculate coding direction after the Go cue (**CD**_**go**_), we averaged *wt* during the first 400 ms of the response epoch (T_go_ < t < T_go_ + 0.4 s). Then we orthogonalized it to **CD**_**delay**_ using the Gram-Schmidt process. To calculate the go direction (**GD**), we subtracted (***r***_lick-right, *t*_ + ***r***_lick-left, *t*_)/2 after the Go cue (T_go_ < t < T_go_ + 0.1 s) from that before the Go cue (T_go_ - 0.1 s < t < T_go_), followed by normalization by its own norm. In Figure S1H, ramping direction (**RD**) was defined as a vector maximally distinguishing the mean activity before the trial onset (0.6 s window) and the mean activity before the Go cue (0.1 s window). All directions were orthogonalized to each other using the Gram-Schmidt process in Figures 1F and S1H. In Figure S6E and S6F, stimulation direction (**SD**) was defined as a vector maximally distinguishing control and stimulation trials after the Go cue/stimulation (T_go_ < t < T_go_ + 0.1 s).

In Figures 2, 6, 7, S1G, S1I-S1K, S2, S6 and S7, we randomly selected 50% of unperturbed correct trials to calculate directions in individual recording sessions. We then projected the spike rate in the remaining trials to the calculated directions as inner products to obtain trajectories. To pool trajectories across sessions, we normalized projections in each session based on the mean trajectories of the unperturbed correct trials. Projection to **CD**_**delay**_ in each session was normalized by the mean activity before the Go cue (T_go_ - 0.1 s < t < T_go_) so that activity in the lick left trial becomes 0 and that in lick-right trial becomes 1. Projection to **CD**_**go**_ in each session was normalized by the mean activity after the Go cue (T_go_ < t < T_go_ + 0.4 s) so that activity in lick left trial becomes 0 and that in lick right trial becomes 1. **GD** in each session was normalized by the mean activity around the Go cue (mean activity difference before and after the Go cue, 100ms window, to be 1). The grand medians are shown in plots. Hierarchical bootstrapping was used to estimate standard errors of median: first, we randomly selected sessions with replacement and, second, randomly select trials within each session with replacement. Neurons with low spike rates (less than 2 spikes per s) were excluded from the analysis. Sessions with less than five cells or without significant selectivity before the Go cue (100 ms window, p>0.05, ranksum test) were excluded from session-based analysis because directions could not be well defined (34 /129 sessions). In Figure 6E and S6L-S6P, sessions with stimulation-triggered licks were analyzed (21 sessions).

In Figures 1, 5, and S1H, we calculated directions by subsampling cells recorded across sessions (i.e., projection is not based on simultaneous recordings). For each cell, we subsampled 20 unperturbed correct lick right and left trials each to define directions. Then, we projected the spike rate in the other 20 trials to these directions as an inner product to obtain trajectories. We excluded cells with less than 40 correct trials per lick direction. To calculate the “selectivity explained” (Figure 1F), we first calculated the total selectivity as the square sum of selectivity across neurons (square sum of n × 1 vector). Then, we calculated the square of selectivity of the projection along each mode at each time point. To calculate the “activity explained” (Figure S1H), we calculated the square sum of the spike rate after subtracting the baseline (mean spike rate before the sample onset; 0.6 s window) across neurons. Similarly, we calculated the square sum of the activity along each direction after subtracting the baseline. In Figures 1F, 5F, and 5G, the mean of correct lick right and left trials is shown for **GD**, and the difference between correct lick right and left trials is shown for **CD**_**delay**_ and **CD**_**go**_. Projections are boxcar filtered (causal, 10 ms window). In Figure 5F, projections are standardized by the activity before the Go cue (T_go_ - 0.1 s < t < T_go_). To calculate the latency to the Go cue (Figures 1G, 5F and 5G), we first identified the first time point in which the projection passes five times of the standard deviation after the Go cue (T_std = 5_). Then, we defined the latency as the last time point in which the projection passes two times of the standard deviation between (T_go_, T_std =5_]. We subsampled 1000 cells with replacement in Figures 1E-1G and S1H. In Figure 5, to match the data size across brain areas, we randomly subsampled 226 cells with replacement to calculate the direction and projection (except for SNr and NLL, where we had 116 and 23 cells, respectively; in these areas, we randomly sampled 116 and 23 cells with replacement). We repeated this subsampling 1000 times.

### Histology

Mice were perfused transcardially with PBS followed by 4% PFA / 0.1 M PB. Brains were post fixed overnight and transferred to 20% sucrose PB before sectioning on a freezing microtome. Coronal 50 μm free-floating sections were processed using standard fluorescent immunohistochemical techniques. All sections were stained with NeuroTrace® 435/455 Blue Fluorescent Nissl Stain (Thermo Fisher Scientific, N21479). The fluorescent label was amplified with immunohistochemical techniques with rabbit anti-RFP (Rockland Immunochemicals, Pottsdown, PA, 600-401-3790) and goat anti-rabbit 555 secondary antibodies (ThermoFisher Scientific, A27039) or chicken anti-GFP (Thermo Fisher Scientific, A10262) and goat anti-chicken 488 (Thermo Fisher Scientific, A11039). Cholinergic neurons were labeled with a mouse monoclonal antibody to ChAT (Sigma, AMAb 91130) and goat anti-mouse 647 secondary antibodies (Thermo Fisher Scientific, A11039).

Slide-mounted sections were imaged on a Zeiss microscope with a Ludl motorized stage controlled with Neurolucida software (MBF Bioscience). Imaging was done with a 10× objective and a Hamamatsu Orca Flash 4 camera. Each coronal section was made up of 80–200 tiles merged with Neurolucida software. The whole-brain image stack was registered to the Allen Institute Common Coordinate Framework (CCF) (Wang et al., 2020) of the mouse brain using NeuroInfo software (MBF Bioscience, Williston, VT) or with a Matlab based script (Mike Economo, Boston University). For cell counting (Figure S4), neurons labeled with AAV_retro_ were detected with a Laplacian of Gaussian algorithm using NeuroInfo software (MBF Bioscience, Williston, VT). Tips of the probe tracks were annotated manually and transformed into the Allen CCF to estimate recording sites (Figure S5H). In Figures 4A and S4D, pixel intensities are normalized: (signal in each pixel – mean signal in cortical areas)/ (99.5 percentile signal in the image – mean signal in cortical areas); since these subcortical areas do not project to the cortex, the cortical signal was used as a baseline for normalization. In Figure 4B, the mean normalized pixel intensities were averaged per nucleus in thal_ALM_.

To define thal_ALM_, we injected WGA-Alexa555 in ALM (Guo et al., 2017). After registering to Allen CCF, 10 μm voxels with retrogradely labeled cells in more than half of cases in the thalamus (n = 3 mice) were defined as thal_ALM_.

For the fluorescent *in situ* hybridization (Figures S4G and S4H), we performed *in situ* hybridization chain reaction (HCR; Molecular Instruments) (Choi et al., 2018) on 150 μm thick coronal sections, following the protocol described in (Nicovich et al., 2019). We probed chat (H1), gad1(H2) and, slc17a6 (vglut2; H3) with amplifiers conjugated with Alexa546 (for H1), Alexa647 (for H2), and Alexa594 (for H3) (See Supplementary Table 4 for probe sequences). Images were acquired with LSM880 confocal microscope (Zeiss) using Plan-Apochromat 40x N.A. 1.3 Oil objective.

### Acute slice recording

AAV5-EF1α-DIO-hChR2 (H134R)-mCherry (University of Pennsylvania Vector Core) was injected at coordinates from Bregma (in mm); AP −4.7, ML 1.2 and −1.2, DV-3.5 in mice (P21-P28). Mice were used for slice recordings 4-8 weeks after the viral surgery. Animals were deeply anesthetized with isoflurane, perfused transcardially with ice-cold cutting solution containing (in mM): 87 NaCl, 50 sucrose, 25 glucose, 25 NaHCO3, 2.5 KCl, 4 MgCl2, 0.5 CaCl2, and 1.2 NaH2PO4, osmolarity 300–310 mOsm/kg) and subsequently decapitated. Brains were rapidly removed, and 300 μm thick coronal brain slices (Leica VT1000S vibratome, Germany) were prepared in chilled cutting solution, after which the slices were transferred to oxygenated (95% O2/5% CO2) artificial cerebrospinal fluid (aCSF in mM; 118 NaCl, 25 NaHCO3, 10 glucose, 2.5 KCl 2.5, 1.2 NaHPO4, 1.3 MgCl2, 2.5 CaCl2). Slices were kept at 33 °C for 30 min and then kept at room temperature for at least 30 minutes prior to recording.

Acute brain slices were transferred to the recording chamber of an upright microscope (Olympus BX50WI, Japan) fitted with a CCD camera (Michigan City, IN), LED system (Olympus pE-2 CoolLED, Japan) with YFP/RFP filter sets, and a Multiclamp 700B amplifier (Molecular Devices, USA). During the recording of VM neurons, slices were perfused with warmed oxygenated ACSF (32 ± 2 °C) through a gravity-fed system connected to an in-line feedback-controlled heater (TC 324B, Warner, USA). Recording pipettes were fabricated by pulling (Narishige PC-10, Japan) borosilicate glass (GC150F, Harvard Apparatus, UK) to a tip resistance of 4–6 MΩ when filled with an internal solution. The internal solution contained (in mM): 135 KMeSO4, 7 NaCl, 10 HEPES, 2 Mg2ATP, 0.3 Na3GTP, and 0.3% biocytin, pH7.3 adjusted with KOH, osmolarity 280-290 mOsm. In experiments where only voltage-clamp and no current-clamp recordings were made, the internal solution contained (in mM): 135 CsMeSO4, 8 NaCl, 10 HEPES, 7 Na-phosphocreatine, 2 Mg2-ATP, 0.3 Na3-GTP, 10 EGTA, 0.1 spermine, and 0.3% biocytin (pH, 7.3 with CsOH; Osmolarity, 280-290 mOsm). Electrode offset potentials were corrected prior to giga-ohm seal formation. ChR2-expressing axon terminals were light-activated with 5 or 100 ms whole-field LED illumination at blue excitation wavelengths (470 nm) at 1.8 mW (0.09 mW for subthreshold stimulation). Liquid junction potentials were not compensated for. The series resistance was typically between 10–30 MΩ and was monitored during the experiments. For voltage-clamp average responses, neurons were clamped at −60 mV, and a minimal of 10 pulses (470nm) were given with 10-second intervals. These light-activated response amplitudes were measured from baseline to peak. Response latencies were measured from the onset of the 5 ms light pulse, and jitter was defined as the standard deviation of the latencies. Data were acquired using AxoGraph X (AxoGraph, Australia), filtered at 10 kHz, and digitized at 20 kHz using an ITC-16 board (InstruTech, USA). Off-line analysis was performed with AxoGraph X.

### Statistics and data

The sample sizes are similar to the sample sizes used in the field. No statistical methods were used to determine sample size. During experiments, trial types were randomly determined by a computer program. During spike sorting, experimenters cannot tell the trial type and therefore were blind to conditions. All comparisons using *signrank* and *ranksum* tests were two-sided. All bootstrapping was done over 1,000 iterations. Data sets will be shared at CRCNS.ORG in the NWB format.

---

**Figure S1.**
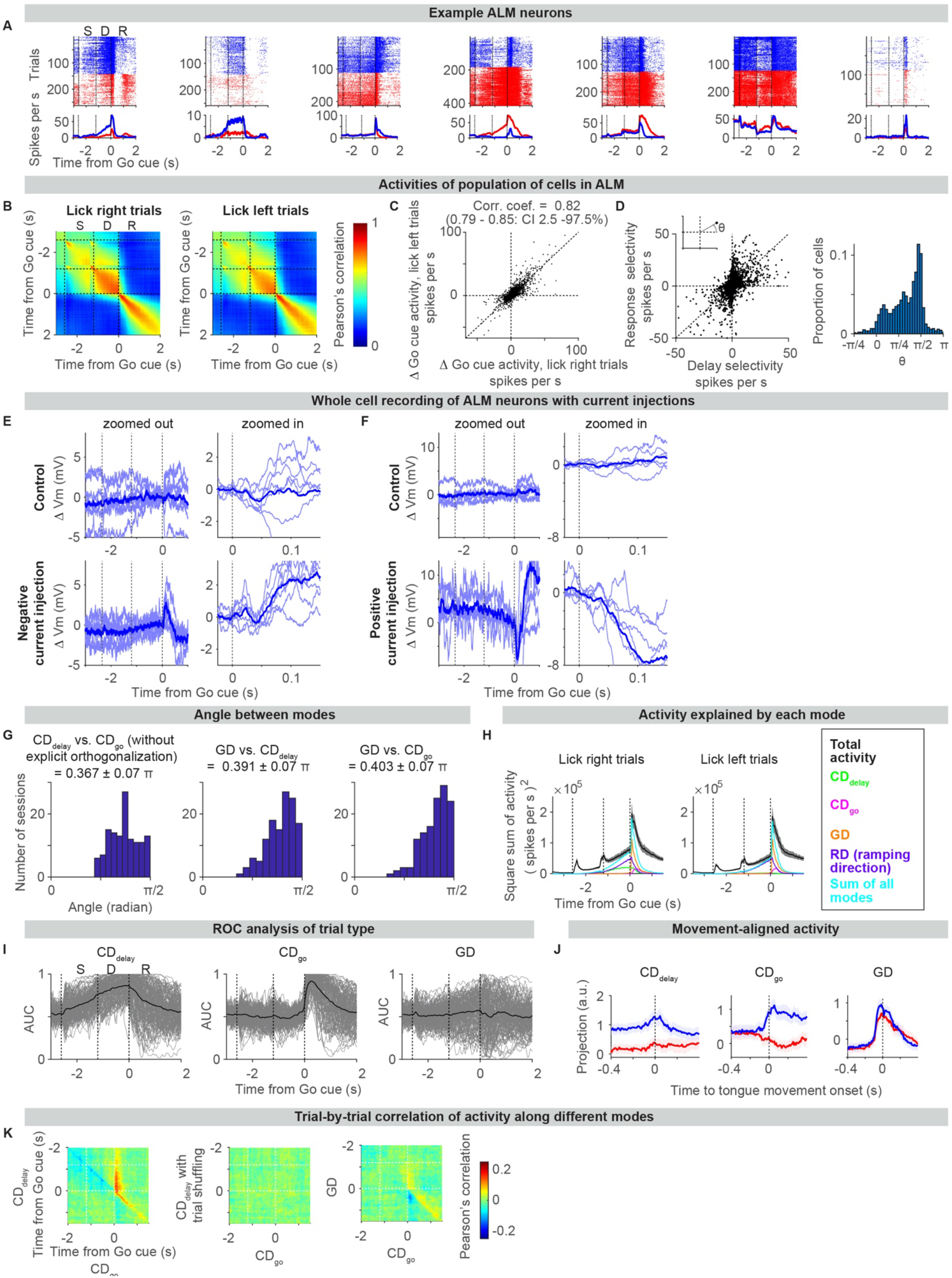
Related to figure 1. Activity underlying motor planning and movement initiation in ALM. A. Example neurons in ALM. Top, spike raster. Bottom, mean spike rate. Blue, correct lick right trials; red, correct lick left trials. Time is aligned to the onset of the Go cue. Dashed lines separate behavioral epochs. S, sample epoch; D, delay epoch; R, response epoch. B. Pearson’s correlation of the population activity vector is low between time points before and after the Go cue. Dashed lines separate behavioral epochs. C. Go cue activity (mean spike rate after the Go cue – mean spike rate before the Go cue; 100 ms window) is similar between lick right (x-axis) and left (y-axis) trials. This is consistent with non-selective **GD**. Circles, individual neurons in ALM (5136 neurons). D. Selectivity during the delay and response epochs is not consistent. Left, the relationship between delay selectivity (mean selectivity during the last 600 ms of the delay epoch) and response selectivity (mean selectivity during the first 400 ms of the response epoch). Circles, individual neurons in ALM (5136 neurons). Inset, the definition of θ (angle in a polar coordinate). Right, histogram of θ across neurons. θ ~ *π*/4 indicates similar selectivity during the delay and response epoch, whereas θ ~ 0 or *π*/2 indicates selectivity is strong only in the delay or response epoch, respectively. E. Increase in conductance of ALM neurons after the Go cue. Membrane potential (Vm) of ALM neurons during the tactile task without (top) or with (bottom) negative current injection in correct lick right trials. Change in Vm (ΔVm) from that before the Go cue (100 ms window) is shown. Thin lines, each neuron. Thick line, mean. Results in correct lick left trials were similar (data not shown). Lower right, the increase in Vm with negative current injections indicates an increase in excitatory current after the Go cue. F. Same as **E** for positive current injection. Lower right, the decrease in Vm with positive current injections indicates an increase in inhibitory current after the Go cue. **E** and **F** imply that increased excitatory and inhibitory currents clamp ALM neurons to different activity levels following the Go cue. G. Histograms of angles between different modes across recording sessions (n = 129 sessions). Left, angles H. between 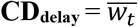 (−0.6 s < t < 0 s; time from Go cue) and 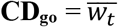 (0 s < t < 0.4 s; time from I. Go cue) are significantly larger than 0 and closer to *π*/2 without explicit orthogonalization, consistent with the inconsistent selectivity before and after the Go cue shown in **D**. Middle and right, **GD** is near orthogonal to **CD**_**delay**_ and **CD**_**go**_. J. The activity in each trial type can be mostly explained by projections to four modes (Methods). K. Decoding of trial type. ROC analysis to distinguish correct lick right vs. left trials using activity along each mode. Thin lines, individual sessions (n = 129 sessions; 50 ms bin). Thick line, mean. L. Activity aligned to movement onset indicates that mode switch happens prior to movement, and is thus not due to efference copy or sensory feedback. Projection of activity along **CD**_**delay**_, **CD**_**go**_, and **GD** aligned to the first time point the tongue was detected based on high-speed videography (e.g., time 100 ms in Figure 1B). M. Trial-by-trial correlation of activity along different modes (mean of 129 sessions; showing correct lick right trials; lick left trials are similar, data not shown). Activity along **CD**_**go**_ after the Go cue (t>0, x-axis) shows high correlation with activity along **CD**_**delay**_ before the Go cue (t<0, y-axis) (left), but not in trial-shuffled controls (middle) or with activity along **GD** (right). High correlation implies that trials with higher activity along **CD**_**delay**_ before the Go cue tend to show high activity along **CD**_**go**_ after the Go cue.

**Figure S2.**
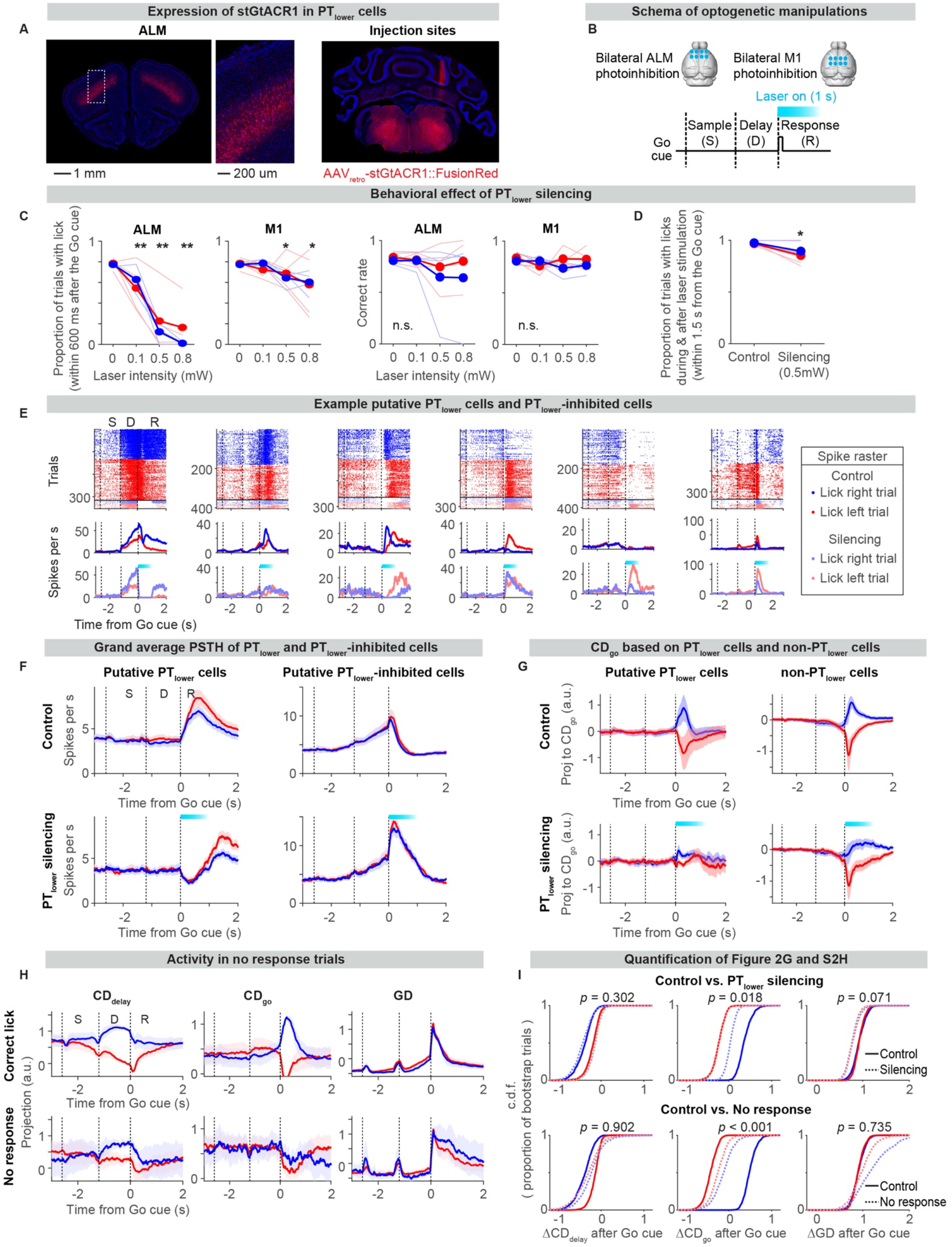
Related to figure 2. ALM motor commands are necessary for movement initiation. A. Coronal brain sections showing bilateral expression of stGtACR1 in PT_lower_ cells. Left, ALM. Right, injection sites in the medulla (see Supplementary Table 1 for coordinates). Red, the fluorescence of FusionRed fused to stGtACR1. Blue, DAPI. B. Schema of PT_lower_ silencing experiments. C. Calibration of laser power. Behavioral effect of PT_lower_ silencing with different laser intensities. Thin lines, individual animals (n = 4 mice). Thick line, mean. PT_lower_ silencing in ALM decreased the proportion of trials with lick without affecting correct rate (probability to lick the correct direction). Because of the significant behavioral effect in ALM with modest effect in M1, we selected 0.5mW for the rest of the experiments. *: *p* < 0.05; **; *p*<0.01 (*Bootstrap* with *Bonferroni* correction for multiple comparisons; null-hypothesis is that the proportion of lick or correct rate in control trials is lower than or equal to those in silencing trials). D. PT_lower_ silencing resulted in loss of lick within 0.6s after the Go cue (Figure 2C), whereas the probability to lick within 1.5 s after the Go cue (laser is on for 1s after the Go cue) was less affected (*p* = 0.003, *bootstrap*). This indicates that licking recovers after the laser stimulation (e.g., Figure 2B). Note that mice lick the correct direction (Figure S2C, correct rate). Same n = 3 mice as in Figure 2C. E. Example putative PT_lower_ cells (cells with a significant decrease in activity during the silencing; left three cells) and putative PT_lower_-inhibited cells (cells with a significant increase in activity during the silencing; right three cells). Top, raster; middle and bottom, mean spike rates of trials with and without PT_lower_ silencing. Blue, mean of all lick right trials (including correct, incorrect, and no lick trials); red, mean of all lick left trials; cyan bar, laser on. F. Grand average PSTH of putative PT_lower_ cells (n = 137 cells), and putative PT_lower_-inhibited cells (n = 111 cells). Note that putative PT_lower_ cells do not have a contralateral bias on average. Line, grand mean of neurons; shading, S.E.M. (*hierarchical bootstrap*); blue, mean of all lick right trials; red, mean of all lick left trials; cyan bar, laser on. G. Projection of activity along **CD**_**go**_ defined only using putative PT_lower_ cells (left) and all putative pyramidal cells excluding putative PT_lower_ cells (right, non-PT_lower_ cells). Consistent with strong silencing of putative PT_lower_ cells regardless of trial types (Figure S2F), activity along **CD**_**go**_ collapsed in both trial types (left). In contrast, **CD**_**go**_ defined by non-PT_lower_ cells showed a reduction in activity only in lick right trials, indicating that contralateral reduction in **CD**_**go**_ is a network effect. Neurons across sessions were pooled for this analysis (n = 137, 737 cells, respectively). Line, grand mean; shading, S.E.M.; cyan bar, laser on. H. Projection of activity along **CD**_**delay**_, **CD**_**go**_, and **GD** in “no response” trials. Activity along **CD**_**go**_ in lick right trials is attenuated. Activity along **CD**_**delay**_ during the delay epoch and **GD** during the response epoch are different from control, yet non-significant within the time window analyzed in **I**. Line, grand median of sessions (n = 24 sessions); shading, S.E.M. (*hierarchical bootstrap*). I. Quantification of change in activity after the Go cue along each direction (activity after the Go cue – activity before the Go cue; 200 ms window). Cumulative distribution function (c.d.f.) across hierarchical bootstrap trials (1000 iterations). *P-value*, hierarchical bootstrap with a null-hypothesis that activity changes in control trials are smaller than or equal to those in silencing (or no response) trials.

**Figure S3.**
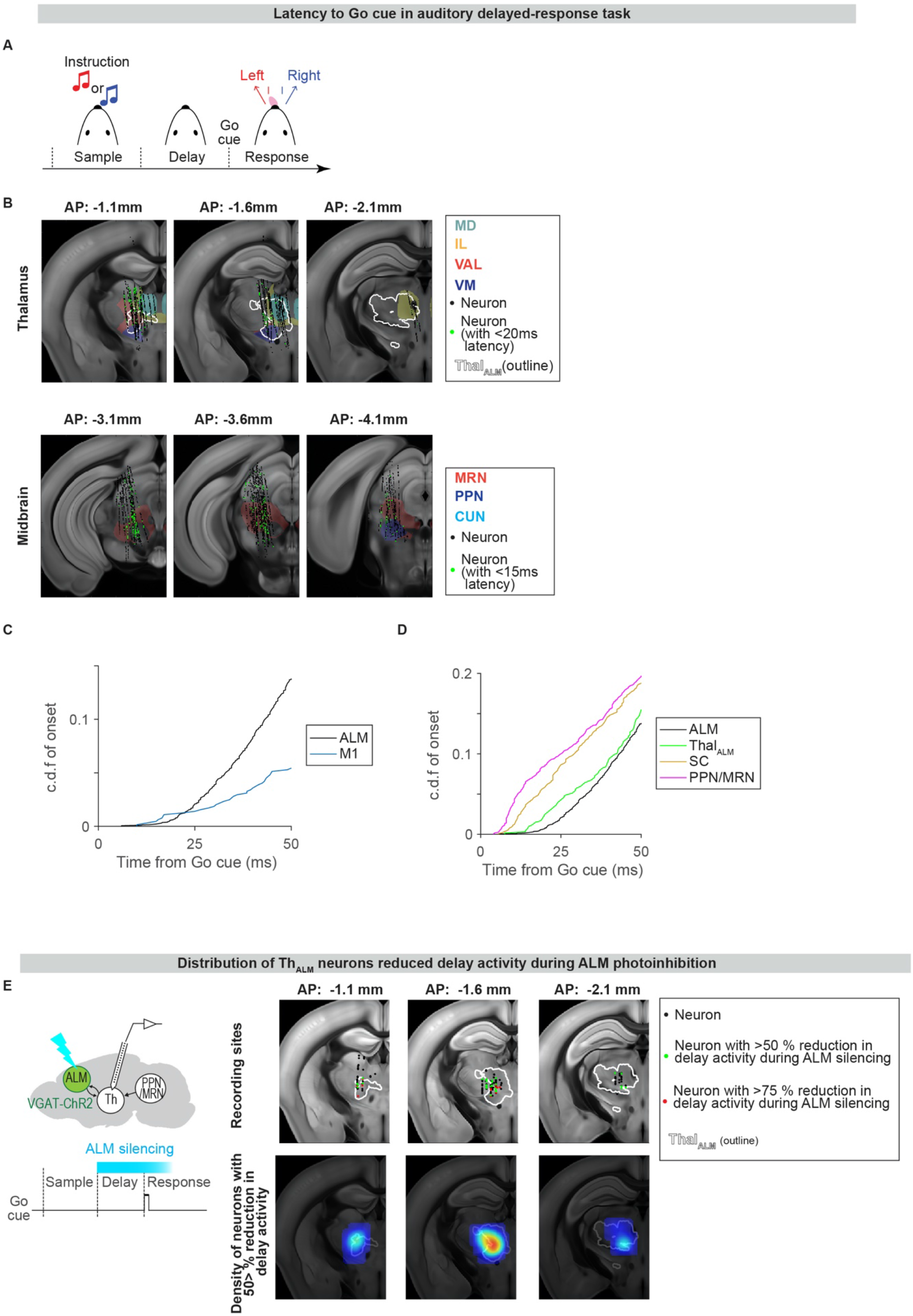
Related to figure 3. Neuronal activity in auditory delayed-response task. Latency to Go cue across brain areas (**B-D**) are similar (latency in PPN/MRN < latency in thal_ALM_ < latency in ALM) in a different task: auditory delayed-response task (**A**). Data in **E** is based on the tactile task (related to Figures 3D-3F). A. Auditory delayed-response task. Tones (3 or 12 kHz) instead of tactile cues were presented during the sample epoch to instruct lick direction. Go cue is 6 kHz FM sound (Methods). B. Recording in the thalamus (top) and midbrain (bottom). Each region filled with color indicates different thalamic or midbrain nuclei. White contour, thal_ALM_. Black dots, location of individual recorded neurons in the Allen common coordinate framework (CCF). Green, neurons with <20 ms (top; in the thalamus) or <15 ms (bottom; in the midbrain) latency to the Go cue. C. Cumulative distribution (c.d.f.) of latency to the Go cue in ALM and M1. Latency (mean ± S.E.M.; time point in which 1% of recorded cells increase activity): 21.1 ± 0.5 ms (ALM; n = 5072 units) and 20.3 ± 4.9 ms (M1; n = 674 units). *p* = 0.402 (bootstrap with a null hypothesis that the latency in M1 is equal to or faster than ALM). D. c.d.f. of latency to the Go cue across brain areas. Latency (mean ± S.E.M.; time point in which 1% of recorded cells increase activity): 21.1 ± 0.5 ms (ALM; n = 5072 units); 16.0 ± 1.5 ms (thal_ALM_; n = 607 units); 10.1 ± 0.8 ms (SC; n = 1145 units); and 7.2 ± 0.5 ms (PPN/MRN; n = 1560 units). E. Distribution of thalamic neurons with decreased delay activity during ALM silencing (left, schema). Note that neurons within thal_ALM_ (white contour) were strongly silenced, consistent with the strong excitatory drive from ALM to thal_ALM_ (Guo et al., 2017). Top, location of individual recorded neurons in the Allen CCF (black). Neurons with more than 50 and 75% reduction in spike rates during ALM silencing (green and red, respectively). Bottom, the density of neurons with more than 50% reduction in spike rates.

**Figure S4.**
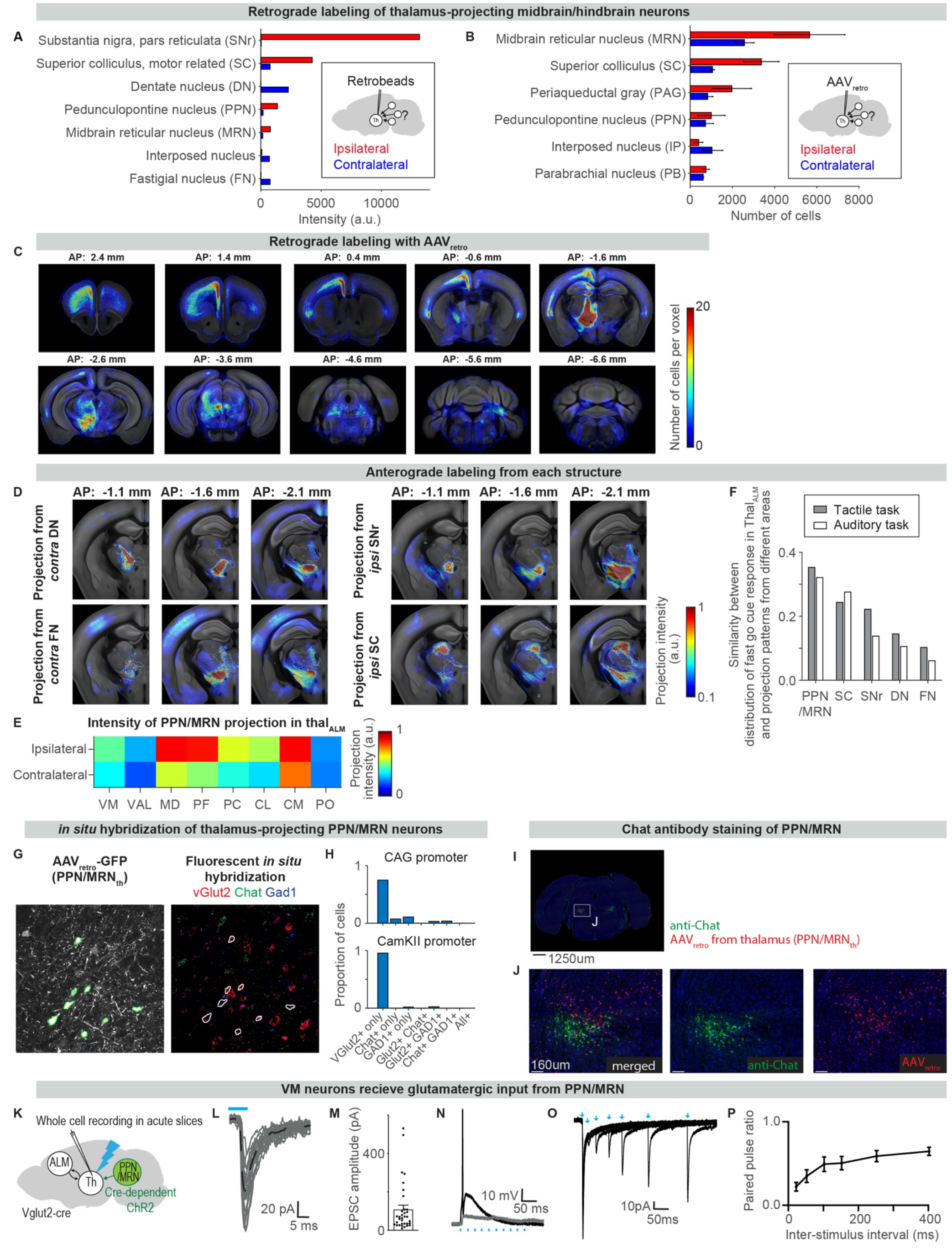
Related to figure 4. Anatomical characterization of neurons projecting to thal_ALM_. Characterization of thal_ALM_-projecting neurons based on retrograde (A-C) and anterograde labeling (D-F). In addition, we confirmed that most thalamus-projecting PPN/MRN neurons are glutamatergic using the fluorescent in situ hybridization (FISH; G, H), immunostaining (I, J), and acute slice recording (K-P). A. Quantification of retrogradely labeled cells in an animal with retrobeads injection in thal_ALM_. The total pixel intensities of retrobeads signal in midbrain/hindbrain areas are shown. Blue, contralateral hemisphere; red, ipsilateral hemisphere to the injection site. Original images of this sample are reported in (Guo et al., 2017). B. Quantification of retrogradely labeled cells in an animal with AAV_retro_ injection in thal_ALM_. The number of labeled cells in midbrain/hindbrain areas are shown. Blue, contralateral hemisphere; red, ipsilateral hemisphere to the injection site. Error bar, standard deviation (n = 2 mice). Some inconsistencies between the retrobeads and AAV_retro_ are caused by known viral tropism (e.g., weak labeling of SNr by AAV_retro_ (Tervo et al., 2016)) and a spread of AAV_retro_ at the injection site beyond thal_ALM_ (Figure S4C). C. Distribution of retrogradely labeled cells in an animal with AAV_retro_ injection in thal_ALM_. Images are registered to Allen common coordinate framework (CCF). AP, relative to Bregma. Heatmap indicates the number of labeled cells per voxel (size: 10 x 10 x 1000 *μ*m). D. Anterograde labeling from distinct subcortical areas to thal_ALM_. Images are registered to Allen CCF. AP, relative to Bregma. Unlike PPN/MRN projection (Figure 4A), projections of these structures are more localized. E. Quantification of anterograde labeling from PPN/MRN to different thalamic nuclei within thal_ALM_. Projection is stronger to the ipsilateral hemisphere. F. Similarity of axonal projection pattern from each subcortical area (i.e., pixel intensities in Figures 4A and S4D), and the distribution of thal_ALM_ neurons with fast go cue responses (i.e., Figure 3C, second row). When ***P*** = pixel intensity and ***D*** = distribution of fast go cue response, we normalized ***P*** and ***D*** by their own norms and calculated the inner dot product between them. A larger number indicates a higher similarity. G. Example images of FISH. Left, PPN/MRN_th_ neurons labeled by AAV_retro_-CamKII-GFP injected in thal_ALM_. Right, FISH of the same section with probes against vGlut2, Chat, and Gad1. H. Quantification of neurotransmitter type (i.e., vGlut2, Chat, and Gad1) of PPN/MRN_th_ cells labeled by AAV_retro_-CamKII-GFP (n = 880 cells) or AAV_retro_-CAG-GFP (n = 404 cells) injected in thal_ALM_. Cells not labeled by any neurotransmitter probes were excluded from the analysis. Cells labeled by CamKII promoter were predominantly vGlut2 positive. I. Anti-Chat immunostaining (green) of a coronal section with PPN/MRN_th_ neurons labeled by AAV_retro_-CAG-H2B::TdTomato injected in thal_ALM_ (red). Consistent with FISH (Figures S4G and S4H), PPN/MRN_th_ were mostly Chat-negative. Blue, Nissl staining. J. Enlarged image of **H**. K. Schema of the acute slice recording experiments. We expressed ChR2 in Glutamatergic PPN/MRN_th_ neurons using AAV-DIO-ChR2-mCherry in Vglut2-IRES-Cre mice. L. Example VM neuron voltage-clamped at −60 mV. Grey lines, twenty individual responses to a single 5 ms 470 nm light pulse (blue bar). Black dashed line, mean. 34/50 VM neurons received PPN input. Latency of EPSC: 2.8 ± 0.8 ms (mean ± standard deviation, n =26 cells). M. Mean EPSC (bar) and individual responses (dots). Error bar, S.E.M. N. Example of evoked action potential (black trace) in response to a 40 Hz 1.8 mW light pulse train (blue bars) at the resting membrane potential, overlaid with potential in response to a subthreshold light pulse train (0.09 mW; greyline). O. Example VM neuron responding to PPN input stimulation with 20, 50, 100, 150, 250, and 400 ms light pulse intervals. P. Average paired-pulse ratio of five VM neurons. Neurons were voltage-clamped at −60 mV. Error bars represent S.E.M.

**Figure S5.**
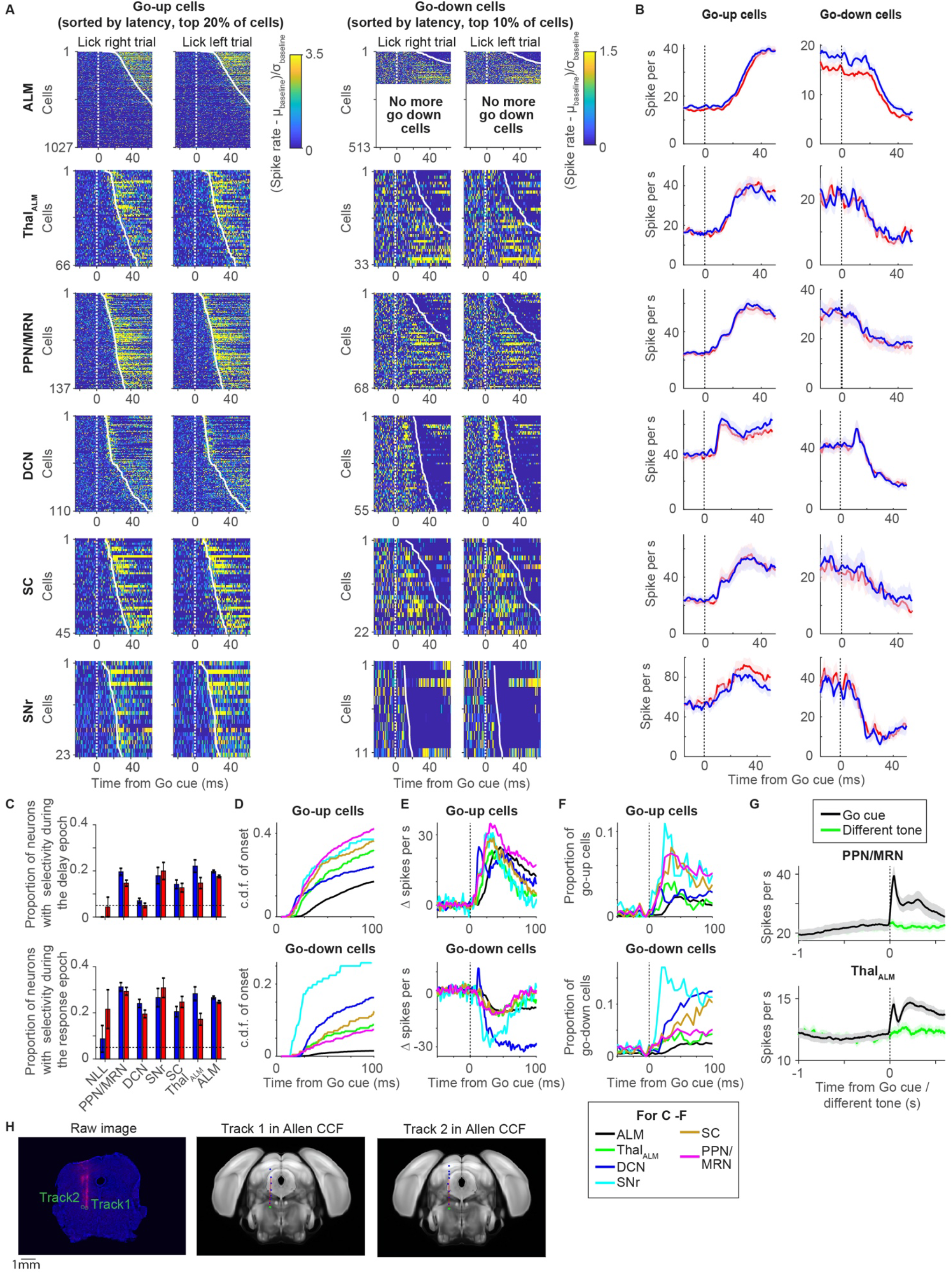
Related to figure 5. Latency to the Go cue across brain areas. A. Spike rates of neurons sorted by their latency to the Go cue in each brain area. From left to right: neurons with an increase in spike rate (go-up cells) in lick right trials and lick left trials and neurons with a decrease in spike rate (go-down cells) in lick right trials and lick left trials. The top 20% of cells and 10% of cells are shown for go-up and go-down cells, respectively. Spike rates were normalized by the spike rate before the Go cue (100 ms) and shown as a heatmap. See Methods for the number of cells recorded in each area. B. Grand average PSTH of the go-up and go-down cells. Line, grand average; shading, S.E.M. (*bootstrap*); blue, lick right trial; red, lick left trial. C. Proportion of neurons with selectivity during the delay (top) or response (bottom) epoch in each area. Blue, lick right trial; red, lick left trial; error bar, S.E.M. (*bootstrap*); dashed line, chance level (*p* = 0.05 as selectivity was defined by ranksum test with α = 0.05). D. Same as Figure 5D, but a broader time-window is shown. Each color indicates a different brain area (box below **F**). E. Overlay of grand average PSTH of the go-up and go-down cells. The mean spike rate before the Go cue (100 ms window) was subtracted. F. Proportion of neurons with significant (ranksum test, *p* < 0.01) increase or decrease in activity after the Go cue (compared to before the Go cue; 100 ms window) at each time point (10 ms bin). G. Grand average PSTH of neurons in PPN/MRN (top) and thal_ALM_ (bottom) in trials with the Go cue (black) or a different tone (green). Related to Figure 5G. In both areas, neurons specifically responded to the Go cue. Both lick right and left trials were pooled. H. Identification of recording location. Probes were painted with CM-DiI, which leaves a track with fluorescent signal. After slicing the brain, each 50 *μ*m thick section was imaged (left, an example section; green circles, tips of the probe). After annotating each track, the images were registered to Allen CCF (Methods). Middle and right, track 1 and 2 in Allen CCF; Red line, estimated probe location in Allen CCF; green dot, estimated tip location; blue dots, other markers placed along the track to estimate the probe location.

**Figure S6.**
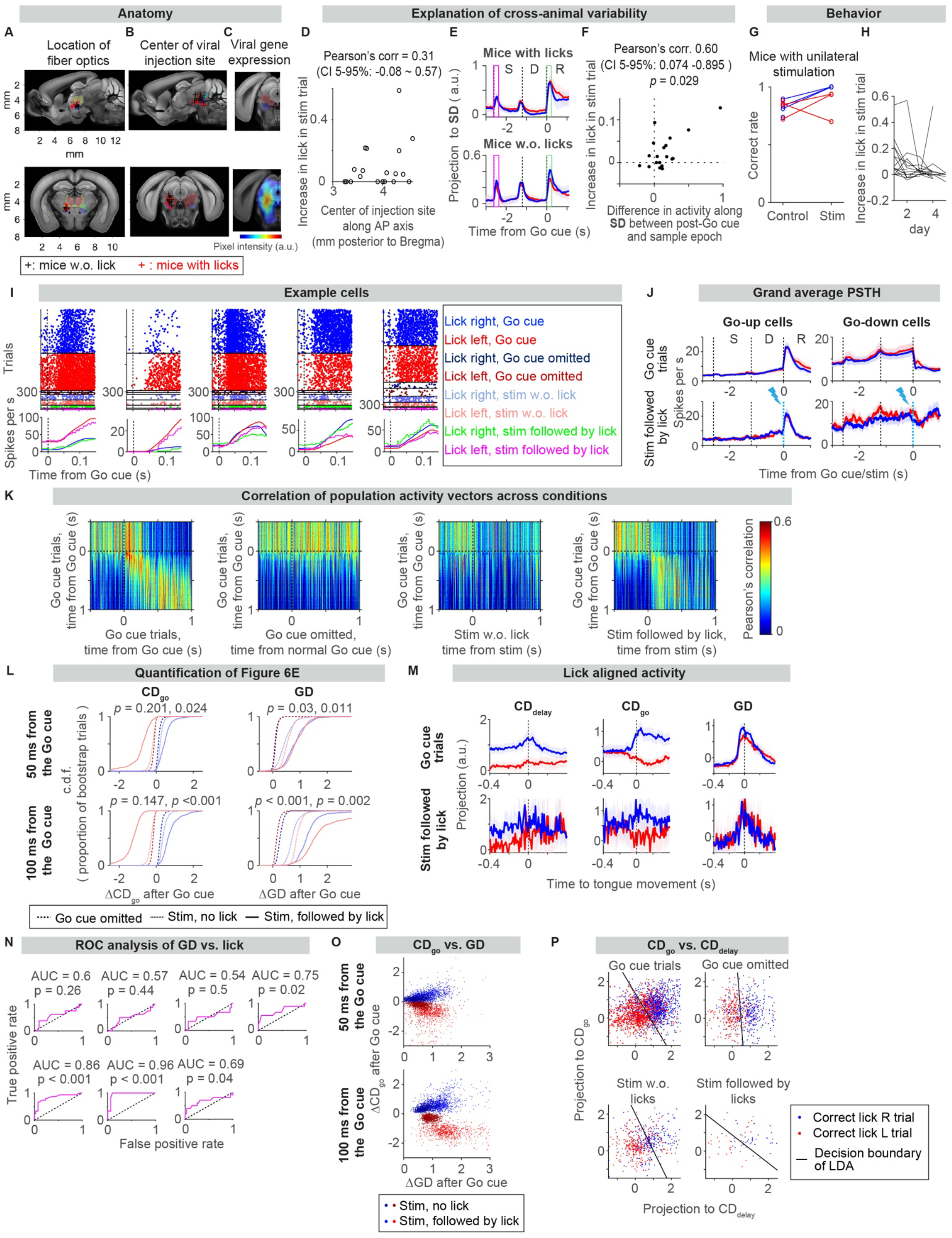
Related to figure 6. Stimulation of thalamus-projecting PPN/MRN triggers lick. A. Anatomical location of fiber optics in the thalamus. Each region filled with a color indicates a different thalamic nucleus. Red, MD; yellow, IL; green, VAL; blue, VM. After recordings, brains were imaged and registered to Allen CCF (n = 20 mice). Black cross, tips of fiber optics in mice without stimulation-triggered lick; Red cross, the same in mice with stimulation-triggered licks. Top, sagittal view; Bottom, coronal view (AP −1.38 mm from Bregma). B. Anatomical location of the center of virus injection in PPN/MRN. Each region filled with a color indicates a different midbrain nucleus. Red, MRN; blue, PPN; green, cuneiform nucleus. Same animals as analyzed in **A**. Top, sagittal view; Bottom, coronal view (AP −3.92 mm from Bregma). C. YFP (conjugated to ChR2) signal around the injection site (mean of 3 mice). Signal intensity is shown in the colormap. The injection site has the strongest signal (red). Weaker signals (cyan) are projections. Top, sagittal view; Bottom, coronal view (AP −4.1 mm from Bregma). D. Anatomical location of viral injection along anterior-posterior (AP) axis and increase in lick in stim trials (probability to lick in stimulation trials – probability to lick in Go cue omitted trials). There is a trend that posterior injection results in a higher probability of stimulation-triggered licks. We see a similar trend with the GtACR experiment as well (Figures S7A-S7B; HI211 and 215 reduced licks with the weaker 0.25mW laser power). CI, confidence interval based on bootstrap. E. Explanation of cross-animal variability in the probability of stimulation-triggered licks based on ALM activity. We defined a stimulation direction (**SD**), which distinguishes activity with or without stimulation in ALM (Methods). Activities in Go cue trials projected along **SD** are different between mice with (top) or without (bottom) stimulation-triggered licks. In mice with stimulation-triggered licks, activity along **SD** increased mostly after the Go cue (green boxes). In mice without stimulation-triggered licks, the activity along **SD** also increased in response to sensory cues during the sample epoch (tactile cue and sound caused by the pole movement; magenta box). Thus, in the latter animals, the stimulation did not induce activity patterns similar to that induced by the Go cue, explaining the lack of lick. This may be due to the differences in the location of viral injection and/or fiber optics (Figures S6A-S6D). Line, median. Shade, S.E.M. F. The difference in activity along **SD** between the post-Go cue (green in Figure S2E) and sample epoch (magenta in Figure S2E) vs. increase in lick rate in PPN/MRN_th_ stimulation trials. *P-value*, bootstrap with a null hypothesis that correlation is smaller than or equal to 0. Circle, each animal (n = 20 mice). G. Mice with unilateral virus injection licked correct direction in response to the PPN/MRN_th_ stimulation. Blue and red, lick right and left trials, respectively (n = 4 mice; one mouse did not lick in response to the stimulation, and the correct rate is not defined). H. The proportion of trials with stimulation-triggered licks decreased over sessions, presumably because we did not reward stimulation-triggered licks. Lines, individual mice. I. Activity of example neurons in ALM. Top, spike raster. Bottom, mean spike rate. Time is aligned to the Go cue (or timing of the normal Go cue/stimulation). Mean spike rate is shown for Go cue trials and stim followed by lick trials. J. Grand average PSTH of neurons with an increase (left; n = 50 cells) or decrease (right; n = 42 cells) in spike rate after the Go cue (by more than two spikes per s), comparing go-cue trials (top) and stim trials with licks (bottom). K. Pearson’s correlation of population activity vector, (***r***_lick-right *t*_ + ***r***_lick-left *t*_)/*2*, across trial types. Population activity patterns were similar between trials with stimulation-triggered licks and cue-triggered licks (4_th_ panel). n = 211 cells across sessions were pooled. The same number of trials were subsampled across trial types. L. Quantification of activity after the Go cue (50 ms, top; 100 ms, bottom) along each direction. Cumulative distribution across bootstrap trials shown in Figure 6E (1000 iterations). *P-value*, from left to right, comparison between Go cue omitted vs. Stim no lick, Go cue omitted vs. Stim followed by lick. The null hypothesis is that the change in activity after the Go cue in Go cue omitted trials is bigger than or equal to that in Stim trials. M. Activity aligned to movement onset indicates that mode switch happens prior to movement in the stimulation trials. Projection of activity along **CD**_**delay**_, **CD**_**go**_, and **GD** aligned to the first time point the tongue was detected based on high-speed videography. The Go cue trials data (top) is a duplicate of Figure S1J, shown here for comparison. The same n = 21 sessions (12 mice) as analyzed in Figure 6E. N. Decoding of stimulation-triggered lick based on activity along **GD**. We performed ROC analysis to test the trial-by-trial relationship between the increase in activity along **GD** after the PPN/MRN_th_ stimulation (100 ms window) and whether the animal licked or not in response to the PPN/MRN_th_ stimulation. We analyzed sessions with more than five trials of stimulation-triggered licks (n = 7 sessions). X-axis, false-positive rate; y-axis, true-positive rate. *P-value*, bootstrap with a null hypothesis that AUC <= 0.5. O. Relationship between the increase in activity after the Go cue along **GD** and **CD**_**go**_ (50 ms, top; 100 ms, bottom). Dot, individual bootstrap trial (1000 iterations). Activity change along **CD**_**go**_ is correlated with that along **GD**. P. Relationship between activity along **CD**_**delay**_ and activity along **CD**_**go**_ (mean activity after the Go cue, time of normal Go cue, or stimulation; 100 ms window). Dot, individual trial (pooled from all sessions with stim-triggered licks; n = 21 sessions); line, decision boundary of Fisher’s linear discriminant analysis (LDA) distinguishing lick right vs. left trials. The vertical line (e.g., in Go cue omitted) indicates no relationship between **CD**_**delay**_ and **CD**_**go**_, whereas negative slopes (e.g., in Go cue trials and stim followed by licks) indicate a strong relationship between **CD**_**delay**_ and **CD**_**go**_.

**Figure S7.**
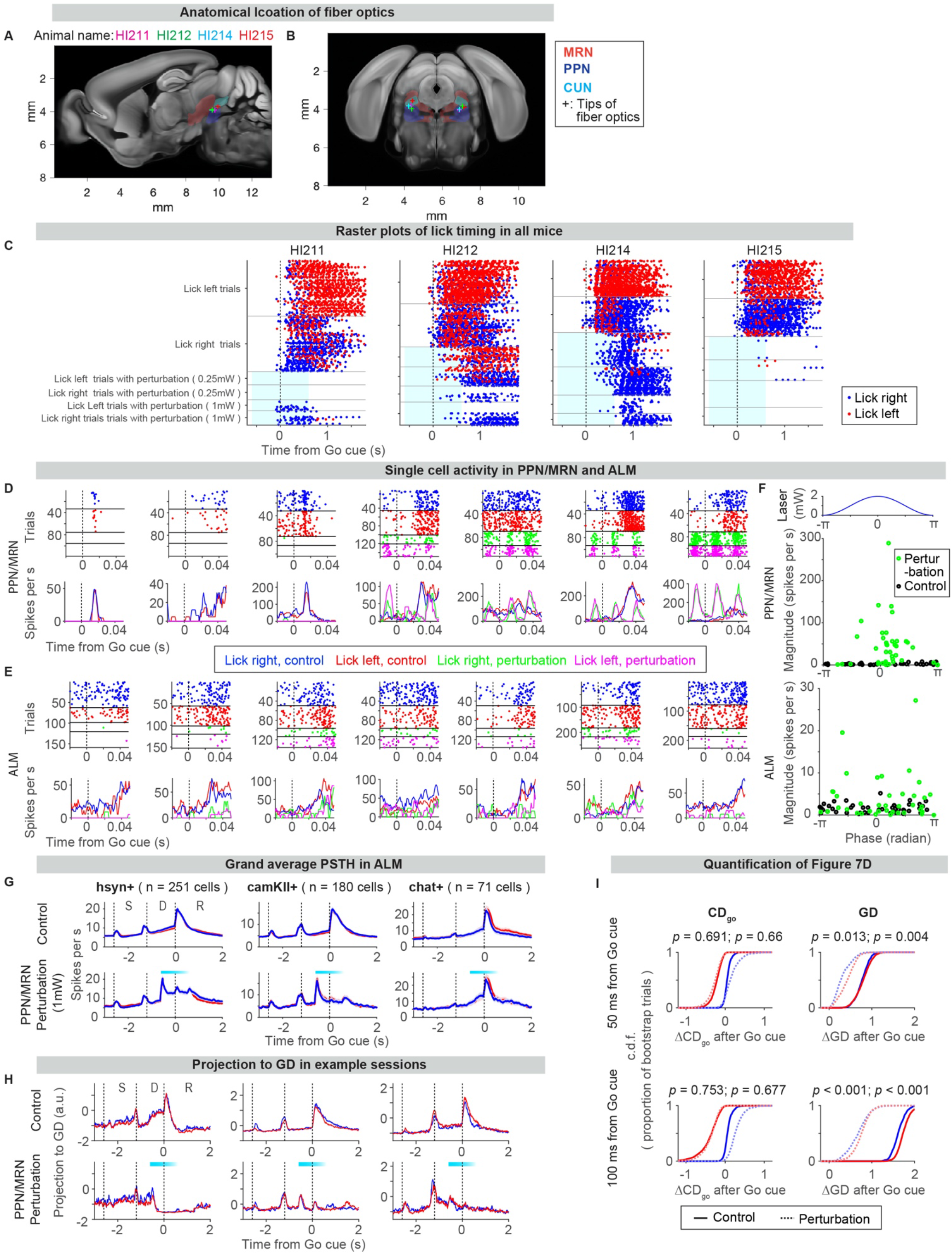
Related to figure 7. Thalamus-projecting PPN/MRN is required for cue-triggered movement initiation. A. Anatomical location of the tips of fiber optics (cross) in PPN/MRN (n = 4 mice; HI## are animal names). Sagittal view. Each region filled with a color indicates a different midbrain nucleus. All data in Figure S7 except **G** is based on animals injected with AAV_retro_-CamKII-Cre in the thalamus. B. Coronal view. Same brains as analyzed in **A**. AP −4.34 mm from Bregma. C. Raster plot of lick timing in all animals. 0.25 and 1 mW indicate laser power used for perturbation. Cyan box, laser on. Behavioral effects were stronger in HI211 and 215. D. Example neurons recorded in PPN/MRN. Top, spike raster. From top to bottom, lick right control, lick left control, lick right with perturbation, and lick left with perturbation trials. Bottom, mean spike rate. Time is aligned to the timing of the Go cue (dotted line). E. Same as **D** for ALM neurons. F. PPN/MRN neurons were modulated by sinusoidal modulation of the laser power. Top, laser intensity in one sinusoidal cycle (25 ms, 40 Hz, mean power: 1 mW). Middle, phase and amplitude of activity of PPM/MRN neurons at 40 Hz (by fast-Fourier transformation, of mean spike activity during the perturbation; 45 cells analyzed in Figure 7C). Circles, individual cells; Black, control trials without perturbation; Green, perturbation trials. Bottom, the same for ALM neurons (44 cells analyzed in Figure 7C). G. Grand average PSTH of ALM neurons in animals expressing GtACR1 in thalamus-projecting hsyn+ PPN/MRN cells (left; from 4 mice), thalamus-projecting CamKII+ PPN/MRN cells (middle; from 4 mice), Chat+ PPN/MRN cells (right; from 2 mice). Top, control trials; bottom, perturbation trials; cyan bar, laser on. H. Example sessions with small increases in activity along **GD** at the laser onset. Top, control trials; bottom, perturbation trials. Cyan bar, laser on. Note that an increase in activity after the Go cue is lost in the perturbation trials. I. Quantification of change in activity after the Go cue (50 ms, top; 100 ms, bottom) along each direction. Cumulative distribution across 1000 bootstrap trials shown in Figure 7D. *P-value*, bootstrap with a null hypothesis that activity change in control trials is smaller than or equal to that in perturbation trials (left *p*-value, lick left trials; right *p*-value, lick right trials).

**Supplementary Table 1.** List of mice and conditions used for optogenetic experiments.

| Figure | Purpose | Mouse genotype, number of mice | Viral injection sites | Photoinhibition method |
| --- | --- | --- | --- | --- |
| Figure 2 | PT<sub>lower</sub> silencing | C57Bl/6J<br>4 mice | AAV<sub>retro</sub>-CamKII-stGtACR1<br>(**Bilateral** injection, coordinate: Bregma AP -6.65mm, ML +/- 1.25mm, DV 4.5mm, 200 nl) | Clear skull prep<br>Photoinhibition 0.5 mW (Bilateral ALM, 8 spots) |
| Figures 3D - 3F | Silencing of ALM during Th recording | PV-IRES-Cre x Ai32<br>2 mice | N/A | Clear skull prep<br>Photoinhibition 1.5mW (Unilateral ALM) |
| Figure 6 | Stimulation of hsyn+ Th-projecting PPN/MRN neurons | C57Bl/6J<br>4 mice | AAV2-hsyn-ChR2-EYFP<br>(**Unilateral** injection, coordinate: Lambda AP +0.2 ~ -0.37mm, ML 1.25mm, DV 2.5mm and/or 3.0mm, 100nl each) | Fiber optics coordinate: Bregma AP -1.5 ~ -1.8mm, ML 0.9mm (**left only**), DV 3.7mm<br>Doric lenses: TFC-200/245-0.37_5mm_TS2.C45 |
|  |  | C57Bl/6J<br>16 mice | AAV2-hsyn-ChR2-EYFP<br>(**Bilateral** injection, coordinate: Lambda AP +0.2 ~ -0.37mm, ML +/- 1.25mm, DV 2.5mm and/or 3.0mm, 100nl each) |  |
| Figure 7 | Perturbation of hsyn+ Th-projecting PPN/MRN neurons | C57Bl/6J<br>4 mice | AAV<sub>retro</sub>-hsyn-Cre<br>(**Bilateral** injection, coordinate: Bregma AP -1.8mm, ML +/-0.9mm, DV 3.9mm, 100nl)<br>+ AAV2/5-hsyn-SIO-stGtACR1-FusionRed<br>(**Bilateral** injection, coordinate: Lambda AP 0.3mm, ML +/-1.25mm, DV 3.0mm, 200nl) | Fiber optics coordinate: Lambda AP 0.325mm, ML +/-1.3mm (**bilateral**), DV 2.7mm<br>Doric lenses: TFC-200/245-0.37_5mm_TS2.C45 |
| Figure 7B | Perturbation of CamKII+ Th-projecting PPN/MRN neurons | C57Bl/6J<br>4 mice | AAV<sub>retro</sub>-CamKII-Cre<br>(**Bilateral** injection, coordinate: Bregma AP -1.8mm, ML +/-0.9mm, DV 3.9mm, 100nl)<br>+ AAV2/5-hsyn-SIO-stGtACR1-FusionRed<br>(**Bilateral** injection, coordinate: Lambda AP 0.3mm, ML +/-1.25mm, DV 3.0mm, 200nl) | Fiber optics coordinate: Lambda AP 0.325mm, ML +/-1.3mm (**bilateral**), DV 2.7mm<br>Doric lenses: TFC-200/245-0.37_5mm_TS2.C45 |
| Figure 7B | Perturbation of Chat+ PPN/MRN neurons | Chat-IRES-Cre<br>2 mice | AAV2/5-hsyn-SIO-stGtACR1-FusionRed<br>(**Bilateral** injection, coordinate: Lambda AP 0.3mm, ML +/-1.25mm, DV 3.0mm, 200nl) | Fiber optics coordinate: Lambda AP 0.325mm, ML +/-1.3mm (**bilateral**), DV 2.7mm<br>Doric lenses: TFC-200/245-0.37_5mm_TS2.C45 |

**Supplementary Table 2.** List of mice used for anatomical experiments. All reagents were introduced into the left hemisphere. After virus injections, expression was allowed for more than two weeks before perfusion.

| Figure | Purpose | Mouse genotype, number of mice | virus or tracer injections |
| --- | --- | --- | --- |
| Figure S4A | Retrograde labeling from Th using Retrobeads | C57Bl/6J<br>2 mice | Red RetroBeads<br>(VM; Bregma AP -1.5, ML 0.85, DV 4.1 mm, 50 nl)<br>Data from Guo et al. 2017 |
| Figures S4B and S4C | Retrograde labeling from Th using AAV<sub>retro</sub> | C57Bl/6J<br>2 mice | AAV<sub>retro</sub>-CAG-H2B::TdTomato<br>(VM left, bregma AP -1.8, ML 0.9, DV 3.9 mm, 100 nl) |
| Figure 4A | Anterograde labeling from PPN/MRN | C57Bl/6J<br>4 mice | AAV2-hsyn-ChR2-EYFP<br>(**Unilateral** injection, coordinate: Lambda AP +0.2 ~ -0.37mm, ML 1.25mm, DV 2.5mm and/or 3.0mm, 100nl each) |
| Figure S4D | Anterograde labeling from other thalamic projecting structures | C57Bl/6J | Data from Gao et al. |
| Figure 4C *1 | Labeling of PPN/MRN neurons | C57Bl/6J<br>2 mice | AAV2/1-CamKII-hChR2::EYFP<br>(PPN left, lambda AP 0.37, ML 1.2, DV 2.5 and 3.0 mm, 100 nl each) |
| Figure 4D *2 | Labeling of Th-projecting PPN/MRN neurons | C57Bl/6J<br>2 mice | AAV<sub>retro</sub>-CamKII-Cre<br>(VM left, bregma AP -1.5, ML 0.9, DV 4.1 mm, 100 nl)<br>+ AAV2/1-hsyn-FLEX-ReaChR::Citrine<br>(PPN left, lambda AP 0.37, ML 1.2, DV 2.5 and 3.0 mm, 100 nl each)<br>+ WGA-Alexa555<br>(ALM, bregma AP 2.5, L 1.5, DV 0.4 and 0.8mm 50nl each) |
| Figure S4G and S4H | HCR | C57Bl/6J<br>2 mice | AAV<sub>retro</sub>-CamKII-GFP or<br>AAV<sub>retro</sub>-CAG-GFP<br>(VM left, bregma AP -1.8, ML 0.9, DV 3.9 mm, 100 nl) |

\*1: AAV2/2-hsyn-ChR2-EYFP resulted in similar results (i.e., strong projection both in thalamus and medulla; data not shown)
\*2: AAV<sub>retro</sub>-hsyn-Cre resulted in similar results (i.e., projection in thalamus without projection in medulla; data not shown).

**Supplementary Table 3.**
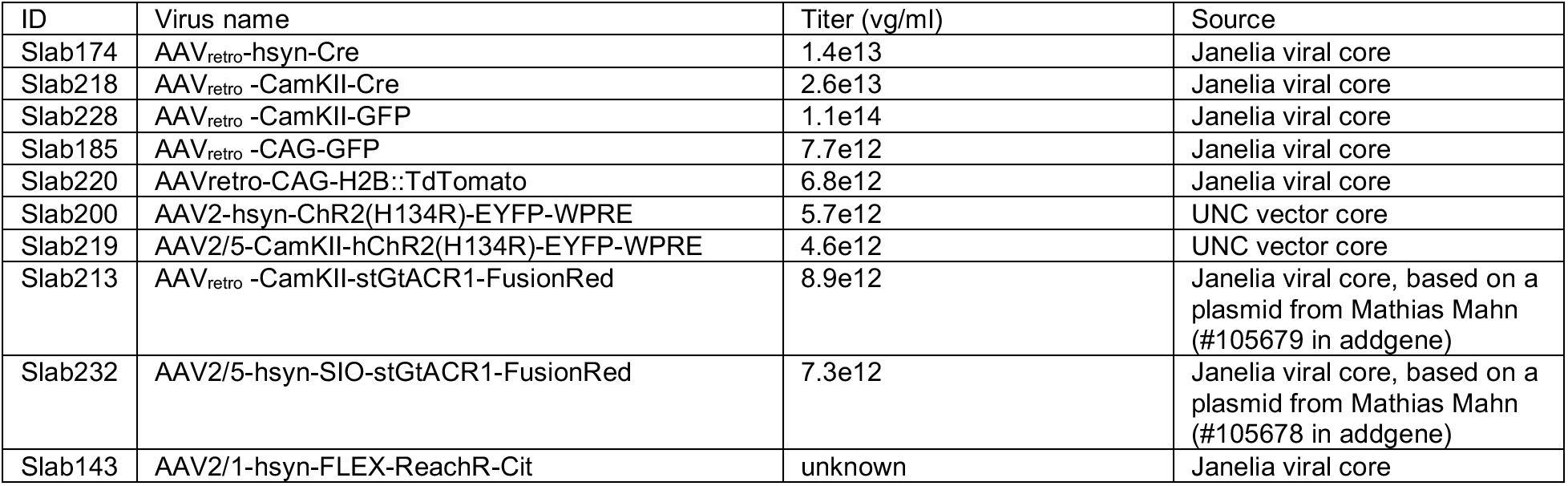
List of viruses used in this paper.

| ID | Virus name | Titer (vg/ml) | Source |
| --- | --- | --- | --- |
| Slab174 | AAV<sub>retro</sub>-hsyn-Cre | 1.4e13 | Janelia viral core |
| Slab218 | AAV<sub>retro</sub>-CamKII-Cre | 2.6e13 | Janelia viral core |
| Slab228 | AAV<sub>retro</sub>-CamKII-GFP | 1.1e14 | Janelia viral core |
| Slab185 | AAV<sub>retro</sub>-CAG-GFP | 7.7e12 | Janelia viral core |
| Slab220 | AAV<sub>retro</sub>-CAG-H2B::TdTomato | 6.8e12 | Janelia viral core |
| Slab200 | AAV2-hsyn-ChR2(H134R)-EYFP-WPRE | 5.7e12 | UNC vector core |
| Slab219 | AAV2/5-CamKII-hChR2(H134R)-EYFP-WPRE | 4.6e12 | UNC vector core |
| Slab213 | AAV<sub>retro</sub>-CamKII-stGtACR1-FusionRed | 8.9e12 | Janelia viral core, based on a plasmid from Mathias Mahn (#105679 in addgene) |
| Slab232 | AAV2/5-hsyn-SIO-stGtACR1-FusionRed | 7.3e12 | Janelia viral core, based on a plasmid from Mathias Mahn (#105678 in addgene) |
| Slab143 | AAV2/1-hsyn-FLEX-ReaChR-Cit | unknown | Janelia viral core |

**Supplementary Table 4.**
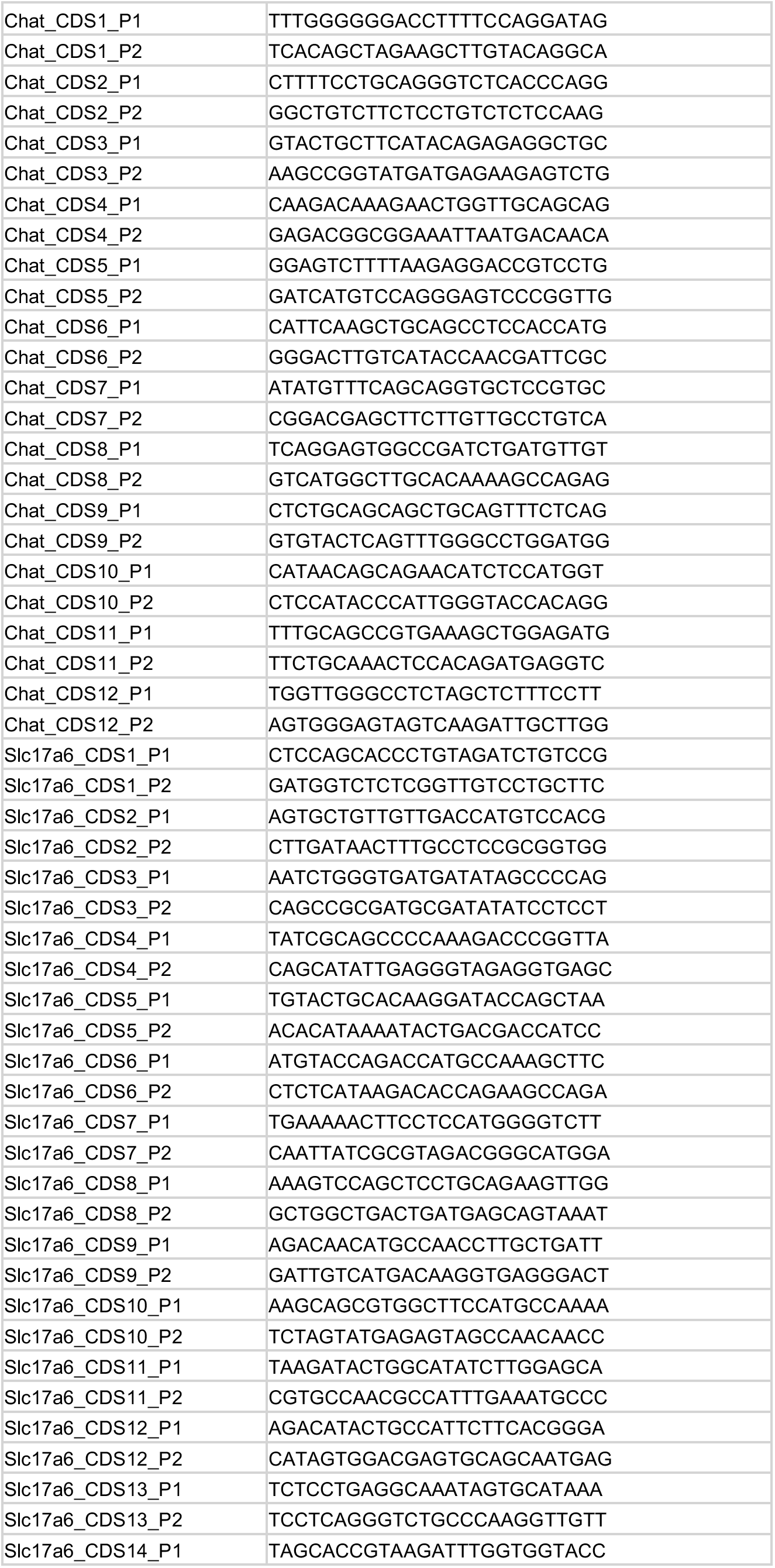
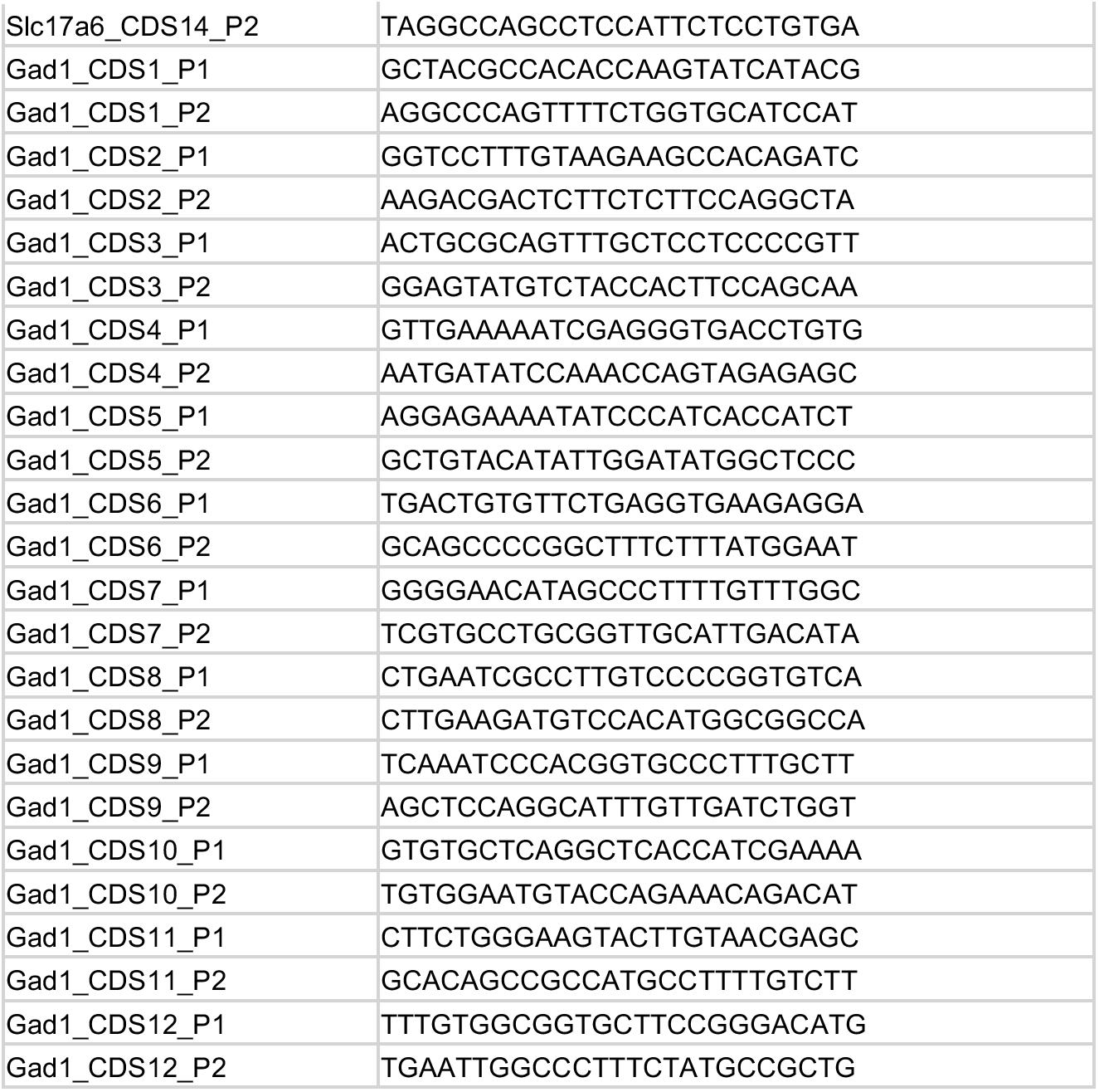
Probe sequences used for HCR. Probes were designed for CDS of *chat*, *slc17a6* and *gad1*.

**Movie S1. Related to figure 1. Example trial with the Go cue**

Top, side view of a mouse with tracking of the jaw (red), nose (blue), and tongue (green). Bottom, movement of jaw and tongue along the dorsoventral axis (a.u.). 400 Hz.

**Movie S2. Related to figure 6. Example trial with thalamus-projecting PPN/MRN stimulation**

Same format as in Movie S1 for a trial with thalamus-projecting PPN/MRN stimulation at time

## Notes

### Competing Interest Statement

The authors have declared no competing interest.

